# The TREM2-R47H Variant Drives Alzheimer’s-Relevant Alterations in Forebrain Organoids Beyond Microglial Populations

**DOI:** 10.64898/2026.02.25.706648

**Authors:** A.S. Kamzina, K.E. Leinenweber, F. Ecca, A. Aldabergenova, M.J. Huentelman

## Abstract

Recent genetic studies highlight microglia as central drivers of Alzheimer’s disease (AD), yet how specific risk variants like *TREM2-R47H* influence broader neurocellular networks remains elusive. Here, we utilize an iPSC-derived forebrain organoid co-culture system to investigate the multi-lineage impact of the *TREM2-R47H* variant. High-resolution transcriptomic profiling, paired with confocal imaging, demonstrate that mutant organoids recapitulate AD-specific pathological signatures. Representative confocal imaging revealed phosphorylated-Tau (pTau) and amyloid-beta (Aβ) internalization by WT microglia, while R47H variants showed a qualitative reduction in pTau accumulation. Single-cell RNA sequencing (scRNA-seq) revealed neurodegenerative transcriptional profiles in *TREM2-R47H* neurons as early as day 139, occurring independently of microglia presence. By day 173, these cell-intrinsic signatures intensified, characterized by disrupted oxidative phosphorylation and impaired maturation trajectories. Interaction analysis further demonstrated that the addition of microglia exacerbated this phenotype; while WT cells adapted to the microglia niche by activating homeostatic, neuro-supportive programs, *TREM2-R47H* cells underwent ‘identity erosion’ and failed to transition into HLA-enriched activation states. This state was characterized by a failure to adopt brain-resident signatures and a divergent shift toward inflammatory myeloid phenotypes. These findings reveal that the *TREM2-R47H* mutation exerts a dual burden: it drives a baseline neurodegenerative state in neural lineages and renders them incapable of proper niche integration. Our study provides an *in vitro* human platform to dissect the interplay between genetic risk and multi-cellular dysfunction, establishing a scalable system for evaluating novel therapeutic interventions and drug screening aimed at restoring neuro-immune homeostasis in AD.

## Introduction

AD is the primary neurodegenerative disease affecting more than 7 million patients in the US and projected to double by 2050^1^ if no preventative or curative solutions are found. Classical hallmarks include amyloid aggregates and neurofibril tangles, however, despite decades of intense research focusing on these protein aggregates, no disease-modifying or long-term efficacy treatments are available. Human genomic heterogeneity has been investigated to explain differences in phenotypes, risk, and age at onset, highlighting more than 80 loci^2^ where *APOE* seems to play a pivotal role together with rare and common variants in *APP*, *PSEN1*, *PSEN2*, and *TREM2* genes^3^, which can lead to familial inheritance forms of AD. However, the ancestry or population specificity of these variants increases the complexity of studying this multifactorial disease and reduces our ability to generalize and finally design new treatments that can ameliorate this dramatic disease. Classical animal models intensely used to decipher Alzheimer’s pathobiology, struggle to recapitulate specific human neurodegenerative features and disease course^4^.

Innovations and advancements in stem cell research, such as development of embryonic stem cells and induced pluripotent stem cells (iPSCs), have enabled the use of human cells *in vitro*^5–7^. Particularly, brain organoids - 3D structures derived from stem cells that self-assemble and somewhat mimic the cellular diversity, function and architecture of the brain - enhanced our ability to study complex brain disorders and provide valuable insights into human biology^8–10^.

Exposure to various combinations of patterning morphogens can yield region-specific organoids such as forebrain, midbrain, hindbrain or spinal cord^11,12^. These brain organoid models contain multiple cell types (neural progenitor cells, astrocytes, neurons) but lack main resident immune cells - microglia that are derived from mesoderm^13^. Currently, there are studies that incorporate microglia with brain organoids to better recapitulate physiological environment in a co-culture system or induce innately^14–16^.

Microglia, the resident immune cells of the CNS, play a crucial role in AD and other neurodegenerative disorders by performing both protective and harmful functions. GWAS studies have shown that many AD risk genes are enriched in microglia^17^, indicating their significant role in AD development. Consequently, numerous studies have explored microglial states in cell models, animal studies, and clinical trials for AD and other neurodegenerative diseases. Recent advances in single-cell technologies have revealed heterogeneous microglial states, including the so-called “disease-associated microglia (DAMs)”, initially described in mouse models of AD^18^ and later reported in human studies as well^19^. These cells have been associated with amyloid plaque clearance and linked to AD pathology. Nonetheless, whether DAMs represent a truly distinct state, and the extent to which such findings translate to human microglia, remains uncertain. Although model systems have been invaluable for uncovering key aspects of microglial ontogeny and function, substantial differences between murine and human microglia - especially in the context of aging and disease - have become increasingly apparent^17,20–22^.

Microglia cells exclusively express triggering receptor expressed on myeloid cells-2 (TREM2)^23^. *TREM2* variants, particularly, *TREM2-R47H* variant increases the risk of developing AD 2-4-fold^24,25^. Evidence for TREM2’s protective role comes from studies showing that *TREM2* knockout or *R47H* variants in AD mice significantly increased tau seeding and spreading around plaques^26^. However, other studies found that *TREM2* deficiency reduced brain atrophy and microglial activation in tau transgenic mice^27^. Notably, TREM2’s impact on amyloid-β and tau pathology appears to differ depending on the disease stage^28^, TREM2 may suppress tau seeding in early AD but exacerbate tau propagation in later stages^29^.

Here, we developed a forebrain-microglia co-culture system combining long-term forebrain organoids (up to day 173 i.e. d173) with isogenic microglia. Using this platform, we show that the *TREM2-R47H* Alzheimer’s risk variant disrupts microglial states and alters other neural cell populations even before microglia are introduced, with neurodegenerative changes detectable as early as day 139 (d139). Notably, the presence of a distinct cluster in WT but not mutant microglia suggest the absence of a potentially protective population in this specific AD variant. Together, these findings provide new insight into how an AD risk gene shapes cellular diversity in the human brain and establish this co-culture system as a physiologically relevant and scalable in-vitro platform for mechanistic studies, disease modeling, and evaluation of therapeutic strategies targeting microglial dysfunction in AD.

## Results

### Generation of human iPSC-derived BOs and microglia

To model a healthy and AD pathological environment, we generated forebrain organoids (BOs) and microglia (Fig. 1a), following the manufacturer’s protocol with minor modifications described in the Methods section. BOs were generated by directed differentiation of *TREM2-R47H* mutant iPSCs and their isogenic WT controls into the dorsal forebrain lineage. Organoids were cultured and collected for scRNA-seq, bulk RNA sequencing and imaging.

**Figure 1.**
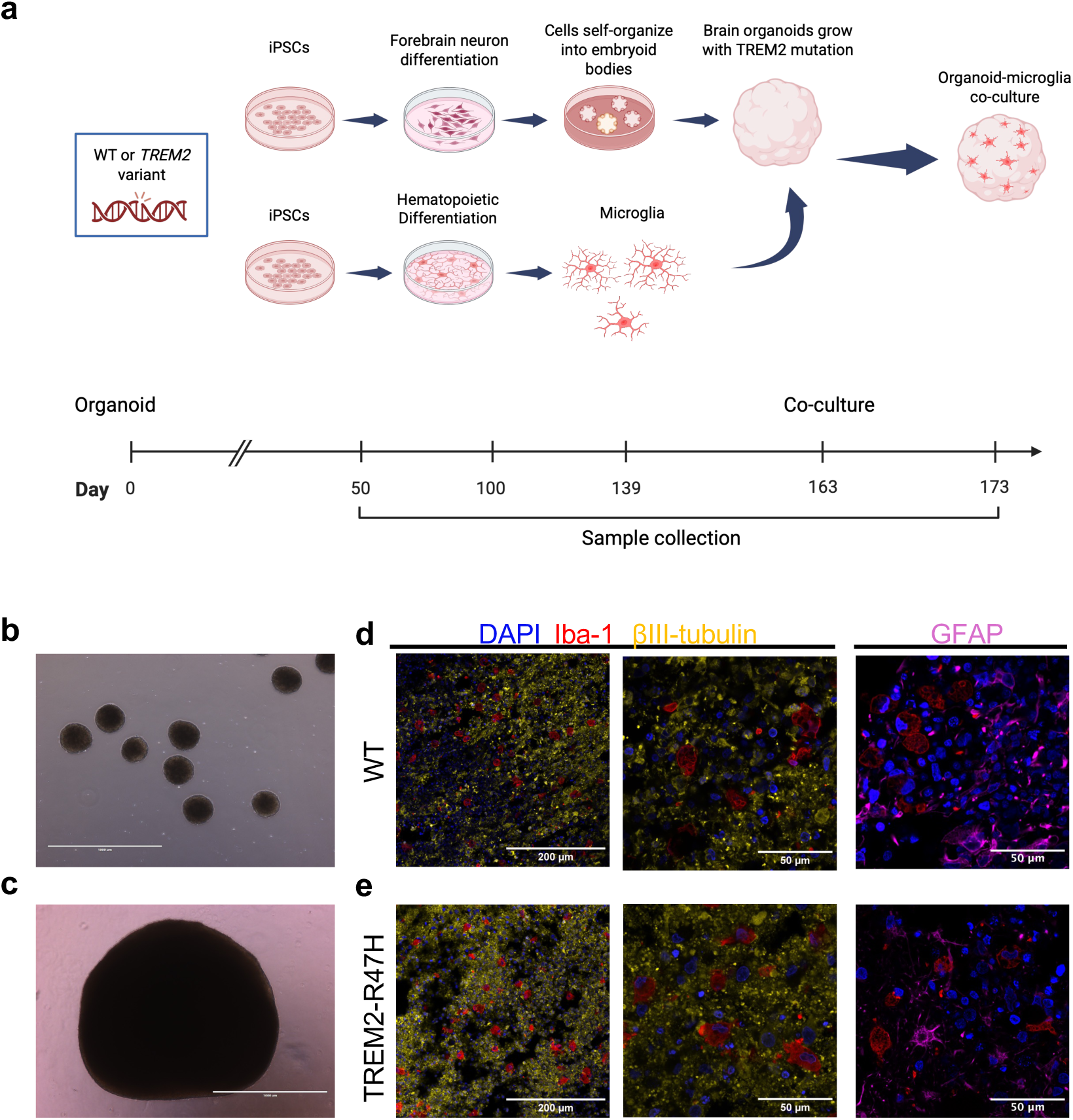
Experimental design and characterization of forebrain organoid-microglia co-culture model. (a) Schematic overview of the experimental workflow. iPSC-derived forebrain organoids and their isogenic microglia counterparts were generated separately and co-cultured on day 163 of organoid development to model AD-related phenotypes *in vitro*. Bottom: experimental timeline indicating differentiation stages and sample collection timepoints. For single-cell RNA sequencing, three to five brain organoids (BOs) were pooled per genotype and processed as one sample. For bulk RNA-seq, three biological replicates per genotype were analyzed, each consisting of three to five pooled organoids. Created in BioRender. Kamzina, A. (https://BioRender.com/q21l969) is licensed under CC BY 4.0. (b) Representative brightfield image of forebrain organoids on day 6. (c) Representative brightfield image of forebrain organoids on day 83. Scale bar, 1000 μm. (d-e) Immunofluorescence staining of organoid sections after 10 days of co-culture showing microglial distribution. (d) WT co-culture. (e) *TREM2-R47H* co-culture. Iba1 (red), βIII-tubulin (yellow), GFAP (magenta) and DAPI (blue). Individual channels were adjusted linearly in Fiji (ImageJ) for brightness and contrast and then merged. Adjustments were applied uniformly across the entire image, and channels were pseudocolored as indicated. Scale bar, 200 μm and 50 μm.

BOs vitality was evaluated via light microscope, based on edge morphology (Fig. 1b, c). Organoids displayed smooth, well-defined surfaces without visible structural disintegration in either WT or *TREM2-R47H* lines across all examined time points. Patterned progenitors preserved their multipotency, giving rise to forebrain neurons as well as GFAPL astrocytes-lineage cells upon terminal differentiation. Immunostaining for β-tubulin revealed preserved tissue architecture without large acellular regions (Fig. 1d, e).

In parallel, we obtained microglia from the same cell lines by inducing iPSCs into hematopoietic progenitor cells (HPCs) followed by differentiation into microglia precursor and mature microglia. Differentiated HPCs exhibited normal circular cells while microglia cells showed typical homeostatic microglial morphology, characterized by circular cell bodies and ramified processes (Supplementary Fig. 1). scRNA sequencing confirmed expression of established microglial marker genes, validating their myeloid lineage identity and confirming the robustness of the method (Supplementary Fig. 2).

To reconstruct the immune-neural interface, mature microglia were introduced into 163-day-old BOs and maintained in co-culture for 10 days. On day 173, BOs were visualized with confocal microscopy with (d173cc10) and without co-culture (d173). The successful integration of microglia into BOs was assessed by immunostaining (Fig. 1d-e), which shows a similar distribution pattern of microglia between the mutated and WT.

To assess possible cellular stress and cell death-associated transcriptional changes, bulk RNA-seq expression of a targeted panel of 10 markers was evaluated across time points for both cell lines beginning at day 100 (Supplementary Fig. 3). Across pre-microglia time points, this gene set did not show pronounced induction. Following microglia co-culture, mutant organoids exhibited increased expression of *HIF1A* and *BAX* at 10 days post-microglia addition, whereas WT organoids showed more modest age-associated changes (Supplementary Fig. 3). When these genes were aggregated into a composite stress score, no significant difference was detected between WT and mutant organoids across matched time points (Supplementary Fig. 4).

Dimensionality reduction using PCA of the bulk RNA-sequencing transcriptomes demonstrated clear separation of samples by developmental timepoint (Fig. 2a). Replicates for each condition tightly clustered together, indicating high consistency across biological replicates. Moreover, within each timepoint, WT and *TREM2-R47H* samples clustered separately after day 100, suggesting a more pronounced genotype-specific transcriptional profile at a later timepoint. The spatial organization of clusters along the PCA axes also implied a temporal trajectory consistent with progressive maturation of the organoids.

**Figure 2.**
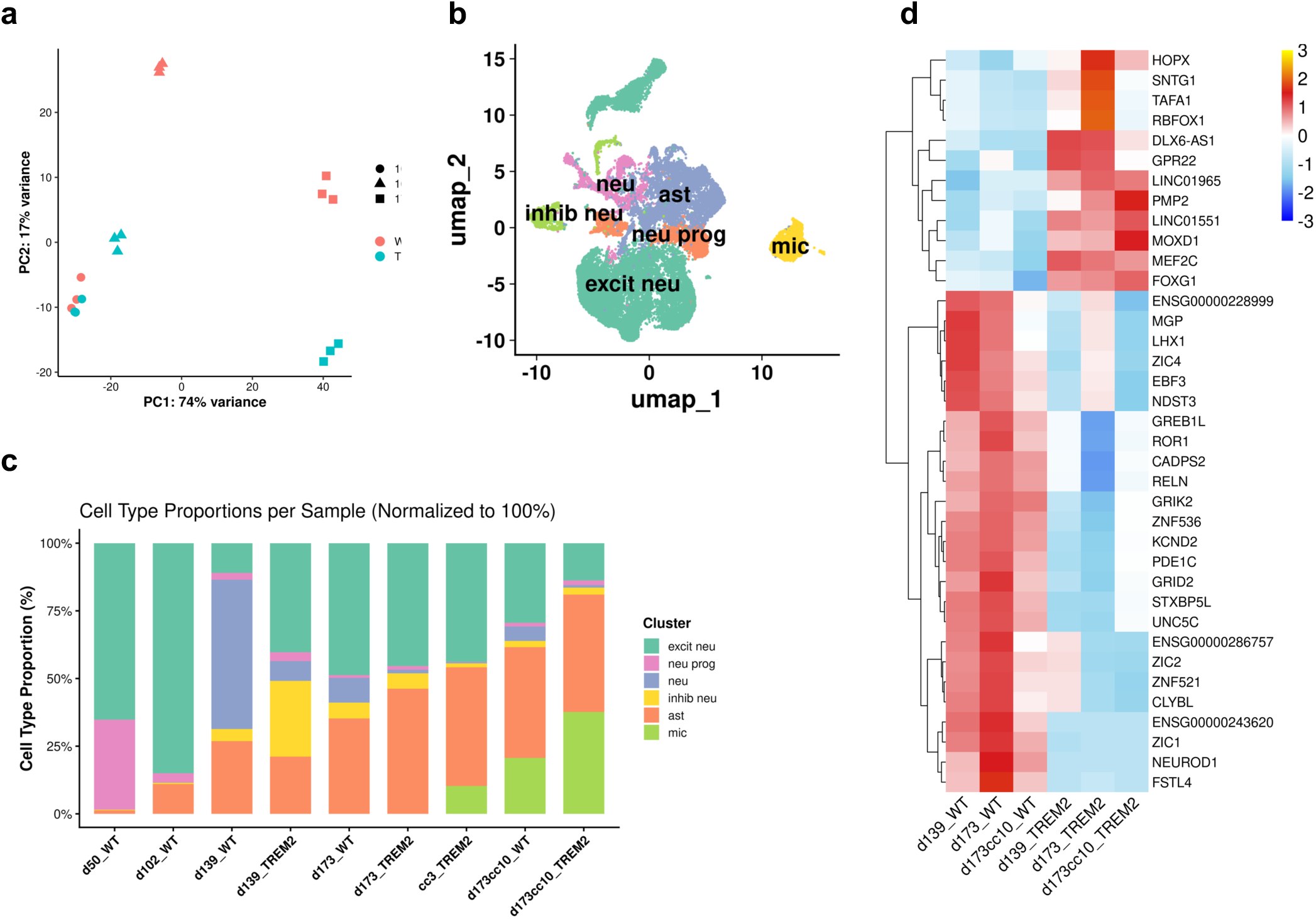
iPSC-derived human BOs have heterogenous cell composition and have low variability between replicates. (a) PCA plot of bulk RNA sequencing after using three replicates with three to five organoids per replicate for days 100, 163 and 173. (b) UMAP and cluster identification of scRNA-seq data from all samples with annotated cell types. (c) Boxplot representing proportions of cell types across the samples with numbers indicated in Table 1. (d) Heatmap of top differentially expressed genes across WT and *TREM2-R47H* groups at d139, d173, and d173cc10. Values represent row-wise Z-scores of average SCT scaled expression across the six plotted groups. Red indicates relatively higher expression and blue indicates relatively lower expression for each gene.

**Table 1.** Cell identity annotated per timepoint and cell line from scRNA.

| Cell type | d50_WT |  | d102_WT |  | d139_WT |  | d139_TREM2 |  | d173_WT |  | d173_TREM2 |  | d173cc3_TREM2 |  | d173cc10_WT |  | d173cc10_TREM2 |  |
| --- | --- | --- | --- | --- | --- | --- | --- | --- | --- | --- | --- | --- | --- | --- | --- | --- | --- | --- |
|  | n | % | n | % | n | % | n | % | n | % | n | % | n | % | n | % | n | % |
| *excitatory neurons* | 4244 | 65.17 | 5251 | 85.02 | 302 | 10.95 | 1125 | 40.26 | 2262 | 48.74 | 688 | 45.38 | 742 | 43.85 | 1296 | 29.41 | 323 | 13.76 |
| *astrocytes* | 83 | 1.27 | 680 | 11.01 | 741 | 26.87 | 591 | 21.15 | 1635 | 35.23 | 701 | 46.24 | 742 | 43.85 | 1805 | 40.96 | 1018 | 43.36 |
| *neuronal progenitors* | 4 | 33.23 | 211 | 3.42 | 70 | 2.54 | 92 | 3.29 | 48 | 1.03 | 21 | 1.39 | 3 | 0.18 | 60 | 1.36 | 41 | 1.75 |
| *microglia* | 2164 | 0.00 | 0 | 0.00 | 0 | 0.00 | 0 | 0.00 | 0 | 0.00 | 0 | 0.00 | 174 | 10.28 | 910 | 20.65 | 884 | 37.65 |
| *neurons* | 0 | 0.06 | 6 | 0.10 | 1521 | 55.15 | 204 | 7.30 | 423 | 9.11 | 20 | 1.32 | 6 | 0.35 | 236 | 5.36 | 22 | 0.94 |
| *inhibitory neurons* | 17 | 0.26 | 28 | 0.45 | 124 | 4.50 | 782 | 27.99 | 273 | 5.88 | 86 | 5.67 | 25 | 1.48 | 100 | 2.27 | 60 | 2.56 |
| *Total* | 6512 | 100 | 6176 | 100 | 2758 | 100 | 2794 | 100 | 4641 | 100 | 1516 | 100 | 1692 | 100 | 4407 | 100 | 2348 | 100 |

### Single-cell RNA sequencing reveals the cellular heterogeneity of organoids derived from WT and TREM2-R47H iPSCs

We examined the cell-specific impact of *TREM-R47H* mutation on the BO transcriptome using the 10X Genomics platform. scRNA-seq was performed on organoids derived from WT and *TREM2-R47H* iPSCs across multiple developmental stages. To capture baseline differentiation trajectories, samples for scRNA-seq at day 50 and day 100 were collected exclusively from WT lines. At later stages (day 139 and 173), both WT and *TREM2-R47H*-derived organoids were profiled to assess the impact of the *TREM2-R47H* variant on cellular composition. At day 173, organoids were analyzed under both microglia-free (d173) and microglia-containing conditions (d173cc10) to evaluate microglial contributions. In addition, a co-culture condition (cc3_TREM2) was included specifically to investigate microglial integration in the TREM2 background at day 3 of co-culture.

Three to five organoids for each time point have been prepared for scRNA-seq. A total of 32,844 cells passed quality control across nine samples (ranging from ∼1,500 to ∼6,500 cells per sample). Correspondingly, total transcript counts (UMIs) per sample ranged from 5.7 million to 25 million, with a mean of about 2,600 – 3,800 UMIs per cell across samples (Supplementary Table 1).

Cells from all timepoints were projected into a two-dimensional space using UMAP based on SCTransform framework (Fig. 2b). Dimensionality reduction and clustering were performed with Seurat, resulting in the identification of ten transcriptionally distinct clusters that were classified into 6 major cell types (resolution = 0.1). Clusters were manually annotated based on established marker genes (Supplementary Fig. 2): neural progenitors (neu prog); neurons (neu); excitatory neurons (excit neu); inhibitory neu (inhib neu); astrocytes (ast); and microglia (mic).

To evaluate the WT cell composition over time and impact of *TREM2-R47H* mutation, we quantified the relative proportion of major cell populations in BOs (Fig. 2c; Table 1). Across all timepoints, WT BOs exhibited a high number of neural progenitor cells on day 50 which decreased in all the later timepoints. Excitatory neurons are dominating inhibitory neurons which is expected since we have dorsal forebrain induction where mainly excitatory neurons develop. Notably, at d173cc10 total neurons constituted 19% of cells in *TREM2-R47H* mutant compared to 38% in WT. This suggests development trajectory and cell fate alteration within the organoid model indicating genotype-dependent shift. To see the difference between cell lines across the timepoint, top 20 DEGs from each of WT and *TREM2-R47H* timepoints (d139, d173 and d173cc10) have been reported in Fig. 2d and full results have been reported in Supplementary Table 2. Notably, the heatmap (Fig. 2d) highlights DE of AD-associated genes, including *RELN*^30^*, MEF2C*^31^*, UNC5C*^32^*, RBFOX1*^33^ and others.

Astrocyte numbers have progressively increased starting when organoids are 100 days old. On the other hand, microglia represented approximately 10% of total cells by day 3 of co-culture, indicating limited initial integration at that timepoint. By day 10, microglial integration increased by at least twofold, with TREM2 mutant organoids exhibiting approximately twice the microglial presence compared to WT controls.

### TREM2-R47H but not WT organoids manifest neurodegeneration

Although TREM2 is almost exclusively expressed in microglia, we hypothesized that TREM2 mutants might impact other cell types, directly or indirectly. To investigate this, we first analyzed RNA expression in d139 organoids with scRNA sequencing. Dimensionality reduction was performed using UMAP, followed by differential expression (DE) analysis, which identified 550 significantly downregulated and 681 upregulated genes (Fig. 3a; Supplementary Table 2). The volcano plot summarizes differential gene expression between TREM2 mutant and WT organoids at day 139, with the 10 most upregulated and 10 most downregulated genes highlighted for visualization (Fig. 3b). KEGG analysis at this timepoint showed reduced enrichment of protein processing and axon guidance pathways and increased enrichment of synaptic signaling pathways in TREM2 mutant organoids relative to WT (Fig. 3c-d; Supplementary Table 3), with GO analyses providing supporting functional context (Supplementary Fig. 5a-c; Supplementary Table 4).

**Figure 3.**
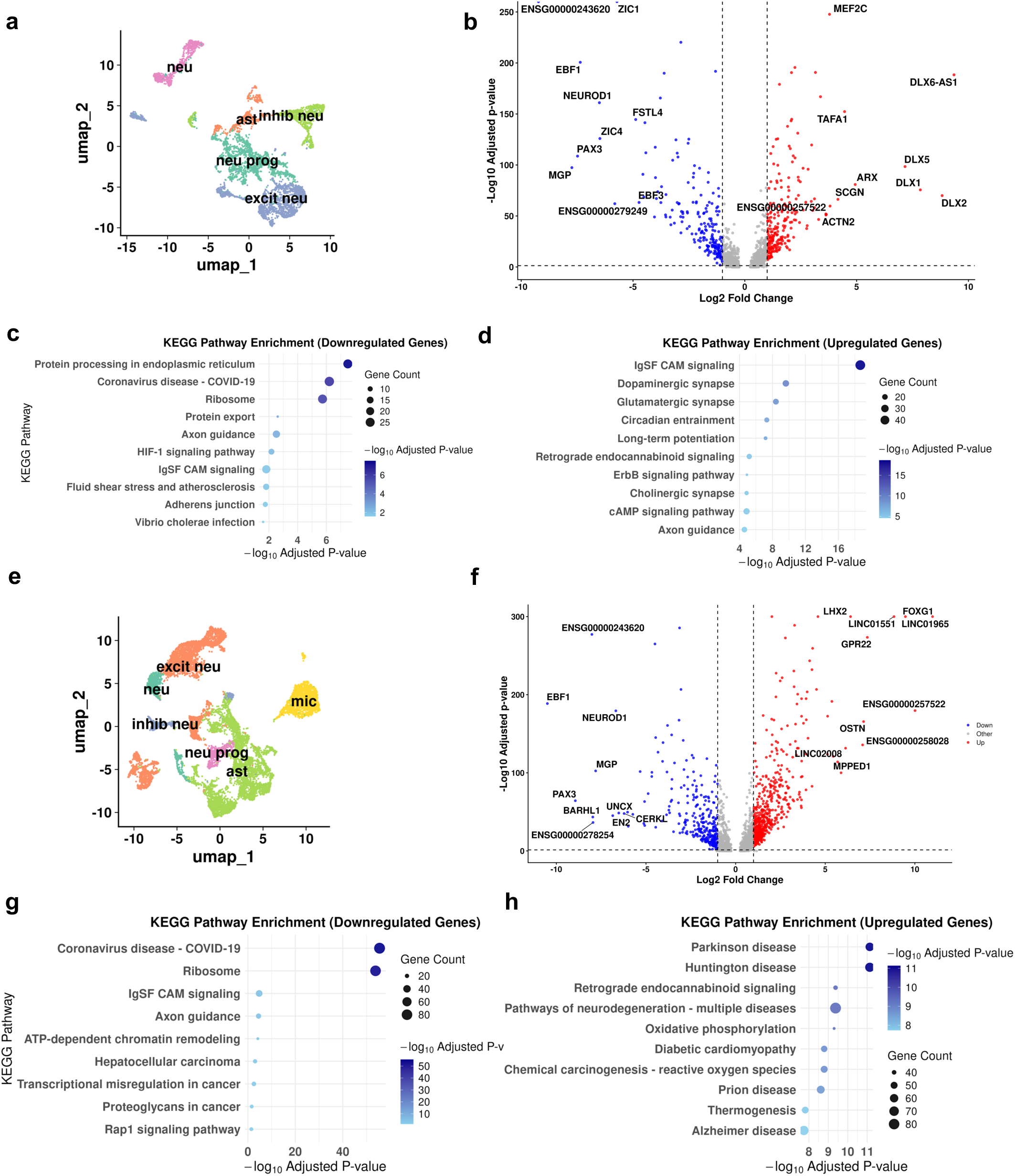
scRNA transcriptomic alterations in WT and *TREM2-R47H* organoids on day 139 and day 173. (a-d) Day 139. (a) UMAP projection of integrated scRNA-seq data with annotated cell types. (b) Volcano plot of differentially expressed genes (|log2FC| > 1, adjusted p < 0.05) comparing *TREM2-R47H* to WT organoids. Positive log2FC values indicate genes upregulated in *TREM2-R47H* relative to WT. (c) KEGG pathway enrichment analysis of genes downregulated in *TREM2-R47H* (i.e., enriched in WT). (d) KEGG pathway enrichment analysis of genes upregulated in *TREM2-R47H*. (e-h) Day 173. (e) UMAP projection of integrated scRNA-seq data with annotated cell types (d173 and d173cc10 samples). (f) Volcano plot of differentially expressed genes (|log2FC| > 1, adjusted p < 0.05) comparing *TREM2-R47H* to WT organoids. (g) KEGG pathway enrichment analysis of genes downregulated in *TREM2-R47H*. (h) KEGG pathway enrichment analysis of genes upregulated in *TREM2-R47H*.

To further investigate the temporal progression of TREM2-associated changes, organoids lacking microglia were collected for analysis on day 173. Using the same UMAP-based analysis pipeline, we identified 775 significantly downregulated and 1,619 upregulated genes, with the 10 most upregulated and 10 most downregulated genes highlighted in the volcano plot (Fig. 3e-f; Supplementary Table 2). KEGG pathway analysis revealed transcriptional differences between *TREM2-R47H* and WT organoids that were primarily associated with neurodegeneration-relevant cellular programs (Fig. 3g-h; Supplementary Table 3). Genes upregulated in *TREM2-R47H* were enriched for pathways related to mitochondrial function and energy metabolism, including oxidative phosphorylation, as well as multiple neurodegenerative disease-associated KEGG signatures (e.g., Alzheimer’s, Parkinson’s, Huntington’s disease, ALS, and prion disease), which share overlapping molecular components linked to mitochondrial activity and cellular stress. In contrast, genes downregulated in *TREM2-R47H* were enriched for pathways associated with axon guidance and glutamatergic synaptic signaling, indicating altered neuron-intrinsic developmental and synaptic programs (Supplementary Table 3). Complementary GO analyses supported these findings by highlighting increased enrichment of mitochondrial and metabolic processes and reduced enrichment of neuronal development and synapse-related categories (Supplementary Fig. 6a-c; Supplementary Table 4).

### Neurons and astrocytes with TREM2-R47H mutation exhibit AD phenotype independently of microglia co-cultures

Initial analysis across all cell types revealed expression of AD-associated markers. Given that microglial dysfunction is a well-established feature of *TREM2* mutations, we sought to determine whether neuronal and astrocytic populations exhibit secondary or cell type-specific transcriptional alterations. To increase resolution in detecting such changes, we performed subclustering restricted to neurons (Fig. 4a, Supplementary Fig. a-c) and astrocytes in microglia-free organoids first at day 139. While KEGG analysis of astrocyte-specific subclusters revealed modest transcriptional differences, neuronal subclusters showed enrichment of neurodegeneration-associated KEGG pathways in both upregulated (Fig. 4b-c) and downregulated (Supplementary Table 5) gene sets, with downregulated terms dominated by mitochondrial and proteostatic modules, and upregulated terms enriched for synaptic signaling and composite neurodegeneration pathways (Supplementary Table 6). GO analysis further supported a shift from translational programs toward neurite- and synapse-associated functions (Supplementary Fig. 7d-f; Supplementary Table 7).

**Figure 4.**
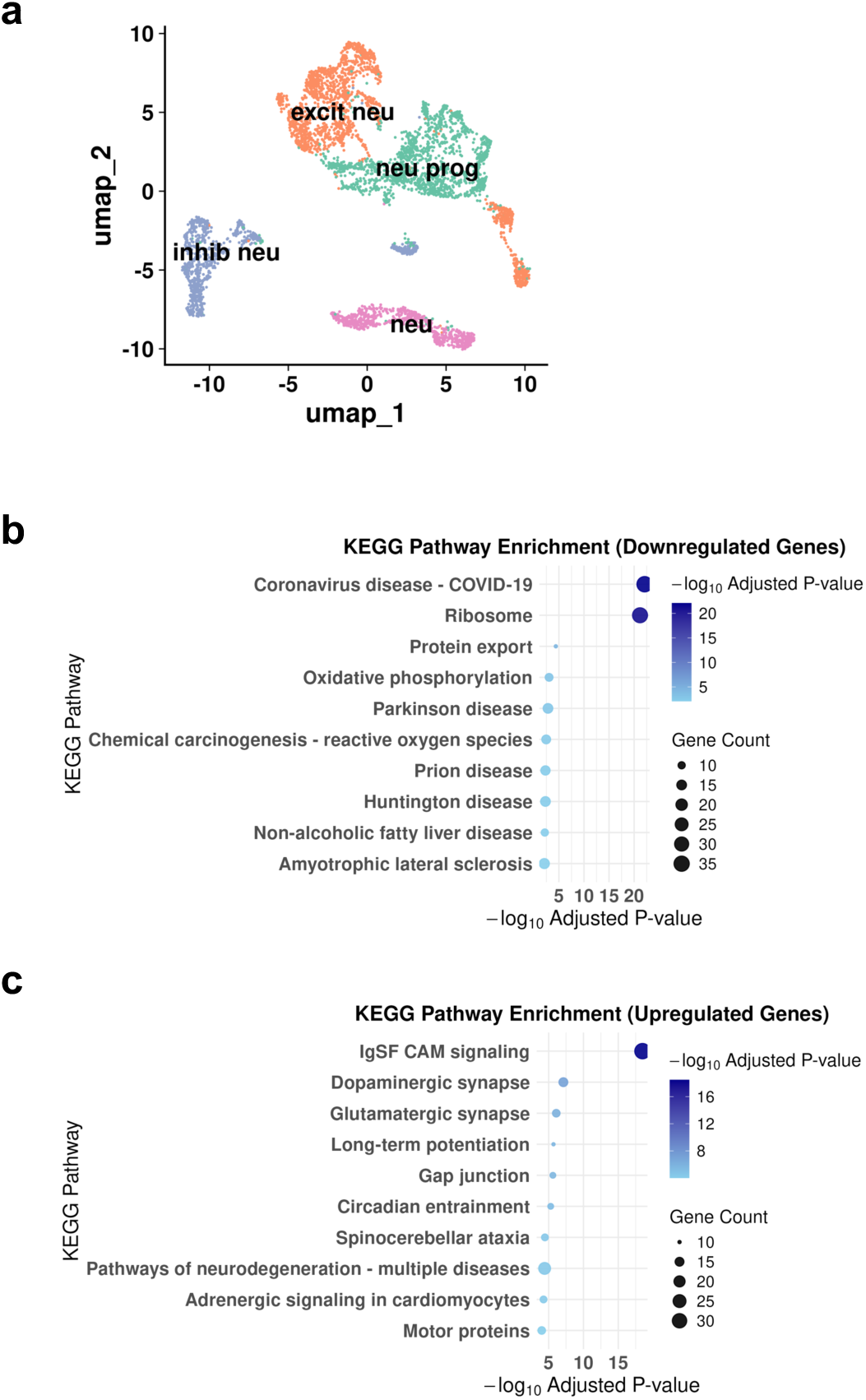
Neuron-specific subclustering and pathway alterations on day 139 organoids. (a) UMAP visualization of neuronal subclustering. (b-c) KEGG pathway enrichment analysis of differentially expressed genes in neurons comparing *TREM2-R47H* to WT organoids.

In d173 organoids, subcluster for inhibitory neurons were depleted in TREM2-R47H (Supplementary Fig. 8b) compared to d139 (Supplementary Fig. 7b). KEGG pathway analysis revealed pronounced transcriptional differences in both neuronal and astrocytic subclusters of TREM2 mutant organoids (Fig. 5b-c) (Supplementary Table 6). In neurons, genes upregulated in *TREM2-R47H* organoids were enriched for multiple neurodegeneration-associated pathways (e.g. Huntington, Parkinson and Alzheimer’s diseases) and synaptic signaling programs, including glutamatergic and GABAergic synapses, whereas downregulated pathways included axon guidance and chromatin remodeling-related processes (Fig. 5b-c). Similarly, in astrocytes, KEGG analysis showed upregulation of oxidative phosphorylation and multiple neurodegeneration-associated pathways (Fig. 5e-f) (Supplementary Table 6).

**Figure 5.**
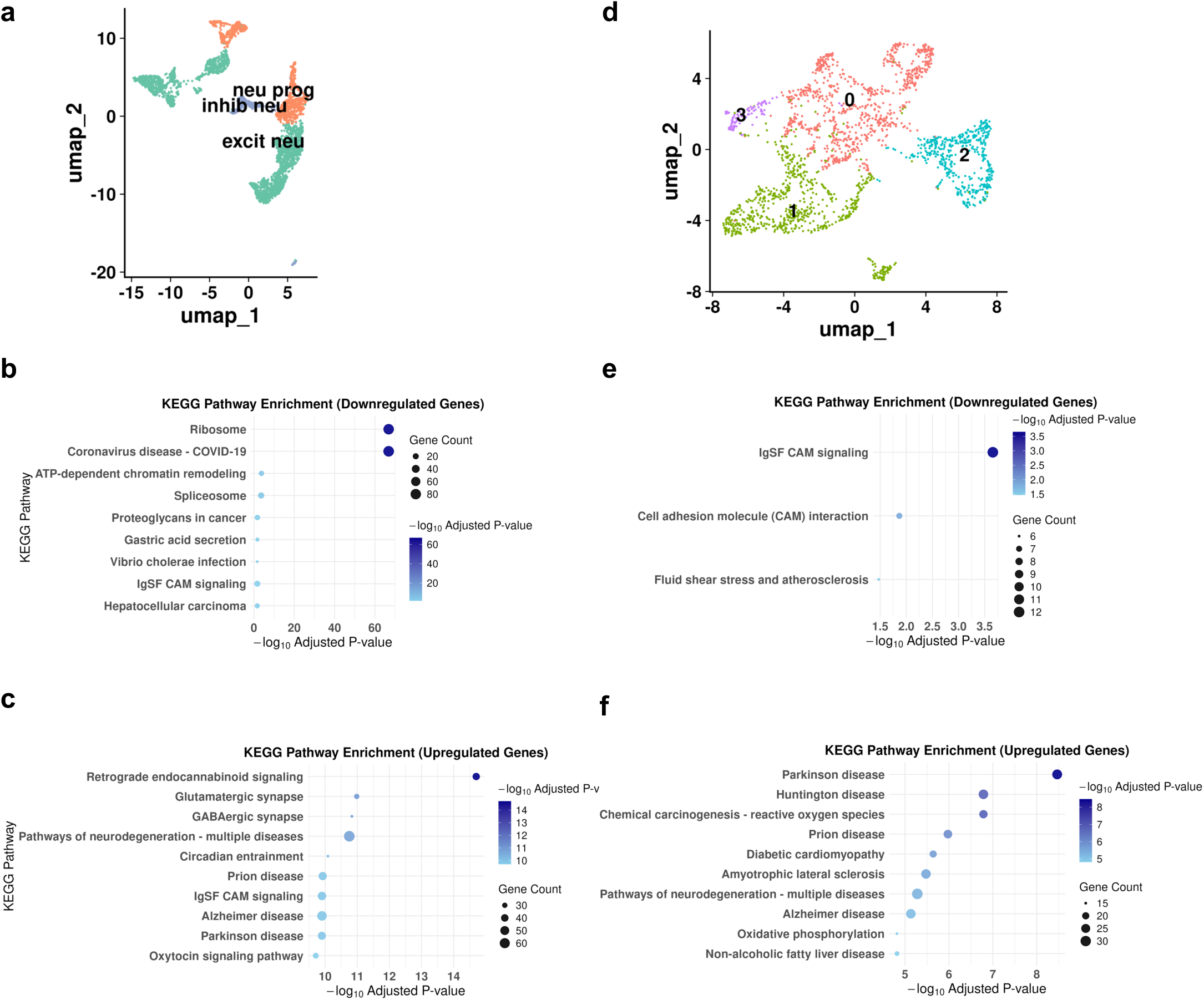
Neuron and astrocyte-specific subclustering and pathway alterations on day 173 organoids. (a) UMAP visualization of neuronal subclustering. (b-c) KEGG pathway enrichment analysis of differentially expressed genes in neurons comparing *TREM2-R47H* to WT organoids. (d) UMAP visualization of astrocytic subclustering. (e-f) KEGG pathway enrichment analysis of differentially expressed genes in astrocytes comparing *TREM2-R47H* to WT organoids.

Together with GO enrichment analyses (Supplementary Fig. 8, 9; Supplementary Table 7, 8), these results indicate that by day 173, *TREM2-R47H* organoids exhibit coordinated, cell-type-intrinsic transcriptional shifts characterized by suppression of developmental and translational programs and increased engagement of metabolic, synaptic, and neurodegeneration-associated pathways.

### TREM2 mutant organoids co-cultured with microglia reveal Alzheimer-specific markers

To visualize AD-related markers and evaluate microglia morphology, we analyzed d173cc10 organoids. To assess colocalization, we did simultaneous staining of microglia (Iba1) and pTau (pTau Ser202, Thr205) which revealed pTau inside WT microglia only and not *TREM2-R47H* (Fig. 6e). Moreover, pTau was sparsely located in the WT BO section. In contrast, in TREM2-mutant BOs, pTau appeared more spread, which may indicate a reduced microglial ability to interact and clear this target (Fig. 6e). Aβ staining (MOAB-2) revealed visually comparable signal between WT and mutant organoids, with no apparent qualitative differences in deposition patterns under the conditions examined (Fig. 6e).

**Figure 6.**
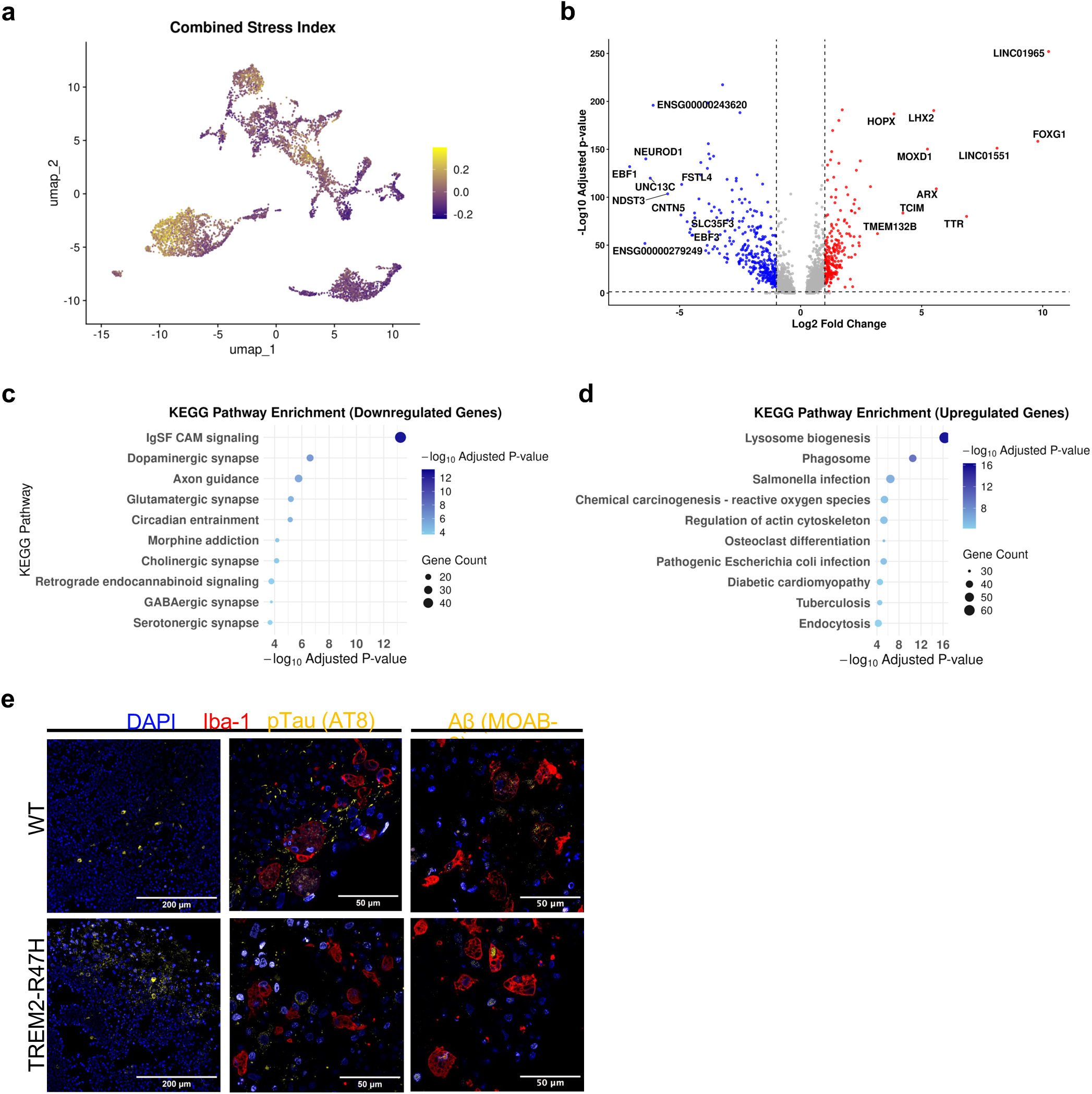
scRNA transcriptomic alterations in WT and *TREM2-R47H* organoids on day 173 co-cultured with microglia for 10 days. (a) UMAP visualization of the combined stress index derived from oxidative stress, ER stress, apoptosis, and microglial activation gene sets; (b) Volcano plot of differentially expressed genes (|log2FC| > 1, adjusted p < 0.05) comparing *TREM2-R47H* to WT organoids. Positive log2FC values indicate genes upregulated in *TREM2-R47H* relative to WT; (c) KEGG pathway enrichment analysis of genes downregulated in *TREM2-R47H* (i.e., enriched in WT); (d) KEGG pathway enrichment analysis of genes upregulated in *TREM2-R47H*; (e) High-resolution immunofluorescence staining of organoid co-culture sections after 10 days of co-culture. pTau (AT8) signal is detected within Iba1⁺ microglia in WT organoid sections (top row), whereas intracellular pTau localization within microglia is markedly reduced or absent in *TREM2-R47H* sections (bottom row). The last column shows Aβ (MOAB-2) staining in organoid sections for both genotypes. Iba1 (red), pTau (AT8) (yellow), Aβ (MOAB-2) (yellow) and DAPI (blue). Individual channels were adjusted linearly in Fiji (ImageJ) for brightness and contrast and then merged. Adjustments were applied uniformly across the entire image, and channels were pseudocolored as indicated. Scale bar, 200 μm and 50 μm.

To investigate the transcriptional impact of microglia co-culture, d173cc10 samples were normalized and analyzed independently to preserve biological signals specific to this experimental condition. scRNA UMAP visualization of the jointly integrated d173 and d173cc10 identified 5 distinct cell types, with the co-culture cells mapping to the same annotated populations (Fig. 3e). We identified *TREM2-R47H*-specific statistically significant DEGs when compared to WT for the same day, including 987 downregulated and 1,172 upregulated genes (Supplementary Table 2). Top 10 up and down regulated genes labeled along with other significant DEGs are visualized in the volcano plot (Fig. 6b). Notably, there is a shift toward upregulation of early developmental and regulatory genes (*LINC01965*, *LHX2*, *HOPX*) while downregulating a large set of genes associated with neuronal differentiation, synaptic function, signal transmission (e.g. *EBF1*, *NEUROD1*, *FSTL4*), potentially indicating altered neuronal maturation and neurotransmission regulators. Full results have been reported in Supplementary Table 2.

KEGG pathway analysis (Fig. 6c-d) revealed a pronounced functional shift in TREM2 mutant organoids where pathways associated with glutamatergic, GABAergic, dopaminergic, cholinergic, and serotonergic synapses, as well as axon guidance, were significantly downregulated (Supplementary Table 3). On the other hand, upregulated genes were enriched for lysosomal, phagocytic, endocytic, and reactive oxygen species pathways, consistent with activated stress and immune responses.

GO enrichment analysis (Supplementary Fig. 10; Supplementary Table 4) of d173cc10 TREM2 mutant BOs compared to WT confirmed a coordinated downregulation of neuronal developmental and connectivity-related programs, consistent with impaired neuronal maturation. In contrast, upregulated GO terms were strongly enriched for regulation of phagocytosis, endocytosis, lysosomal functions as well as amyloid-beta binding pathways.

To assess stress-related transcriptional responses across cell types in the co-culture model, we examined curated gene sets associated with oxidative stress, ER stress, apoptosis and microglial activation. We observed that oxidative stress genes (Supplementary Table 9) are mainly upregulated in microglial and astrocytic clusters (Fig. 6a). Dissociation stress markers^34^ were not found to be increased in these samples. Canonical ER stress and apoptosis-related stress genes in this co-culture model are observed mainly in inhibitory neuron clusters and some astrocytes. Microglia-related genes are highly upregulated in our microglia cluster. A full list of genes used in the analysis have been reported in Supplementary Table 9.

### Astrocytic and neuronal subclusters in co-culture model demonstrate reactive phenotype and classical Alzheimer’s molecular phenotype

In d173cc10 organoids, *TREM2-R47H* astrocytes exhibited modest downregulated pathway enrichment, but showed upregulation of transcriptional signatures consistent with stress and inflammatory responses (eg. wound healing, regulation of apoptosis response to toxic substance in GO and neuroinflammation-like enrichment in KEGG) (Fig. 7b-c; Supplementary Fig. 11, Supplementary Table 6 and 8). While neuronal subclusters (Fig. 7d) displayed depletion of inhibitory neuron sublcuster (Supplementary Fig. 12a), consistent with d173 (Supplementary Fig. 8b), it also showed significant dysregulation of AD associated genes, including reduced expression of *RELN*^30^ and *ERBB4*^35^, together with increased expression of the proteostasis regulator *UCHL1*^36^ (Supplementary Table 7). In line with this, KEGG pathway analysis revealed suppression of key neuroprotective and synaptic signaling pathways, including *ERBB* signaling, as well as multiple synaptic neurotransmission pathways (Fig. 7e) (Supplementary Table 6). In contrast, pathways associated with neurodegenerative disorders such as AD, Parkinson’s, Huntington’s and prion disease were significantly enriched, indicating convergence toward a shared neurodegenerative transcriptional program (Fig. 7f) (Supplementary Table 6).

**Figure 7.**
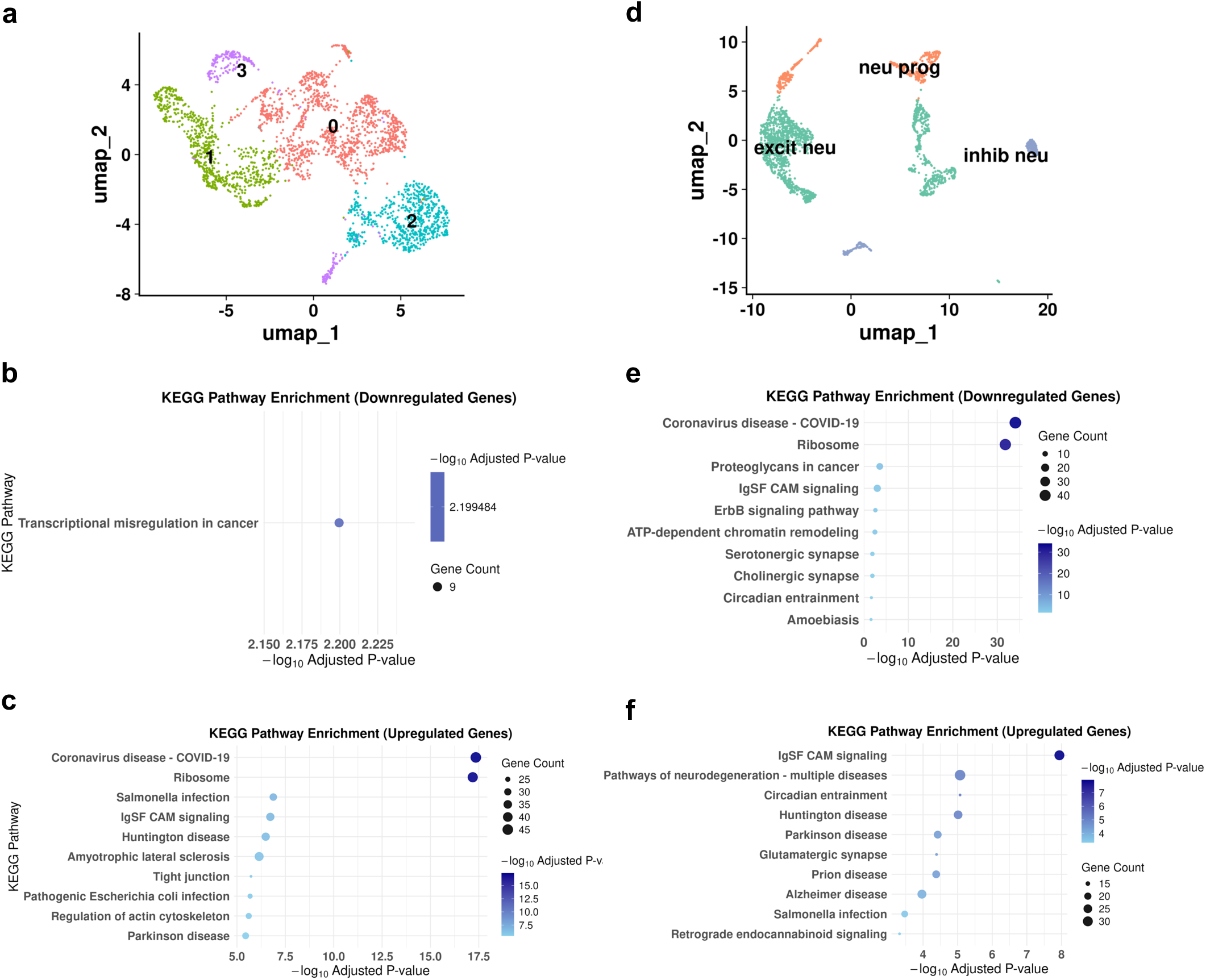
Astrocyte and neuron-specific subclustering and pathway alterations on day 173 co-cultured with microglia for 10 days. (a) UMAP visualization of astrocytic subclustering. (b-c) KEGG pathway enrichment analysis of differentially expressed genes in astrocytes comparing *TREM2-R47H* to WT organoids. (d) UMAP visualization of neuronal subclustering. (e-f) KEGG pathway enrichment analysis of differentially expressed genes in neurons comparing *TREM2-R47H* to WT organoids.

GO analysis further confirmed a marked downregulation of neurodevelopmental and differentiation processes, and upregulation of genes involved in synaptic and postsynaptic organization (Supplementary Fig. 12b-d). Together, these changes define a transcriptional state characterized by synaptic hyperactivity and metabolic stress, consistent with neuronal vulnerability and early neurodegenerative processes.

### Single-cell RNA-seq identifies HLA-enriched cells in WT microglia subcluster

To better differentiate microglial subclusters between conditions, we performed DE analysis on microglia isolated from d173cc10 WT and *TREM2-R47H* organoids (Supplementary table 5).

UMAP-based subclustering identified three transcriptionally distinct microglial subclusters (subclusters 0, 1, and 2) (Fig. 8a). The top 20 DEGs between WT and *TREM2-R47H* microglia are shown in a dot plot (Fig. 8b), and transcriptional differences across microglial subclusters are summarized in the heatmap (Fig. 8c). Overall, the mutant microglial subcluster showed downregulation of multiple HLA-related genes and *CD74*, alongside upregulation of *APOC1* and *CCL3*, genes associated with AD pathology and neuroinflammation^37,38^. Cluster 0 showed high expression of *TREM2, TMEM176A* and *S100A9*, which are commonly associated with disease-associated microglial states described in neurodegenerative contexts (Fig. 8c). While microglial subclusters 0 and 1 were present in both genotypes, subcluster 2 was detected exclusively in WT organoids (Supplementary Fig. 13a). This WT-specific population represented a relatively small fraction of the total microglial compartment (82 cells, approximately 9% of WT microglia) yet exhibited a clearly distinct transcriptional profile (Fig. 8c) characterized by high expression of antigen presentation-associated genes, including *CD207, CD1A,* and *HLA-DQA2,* together with additional MHC class II-related transcripts (Fig. 8c; Supplementary Table 10 and 11).

**Figure 8.**
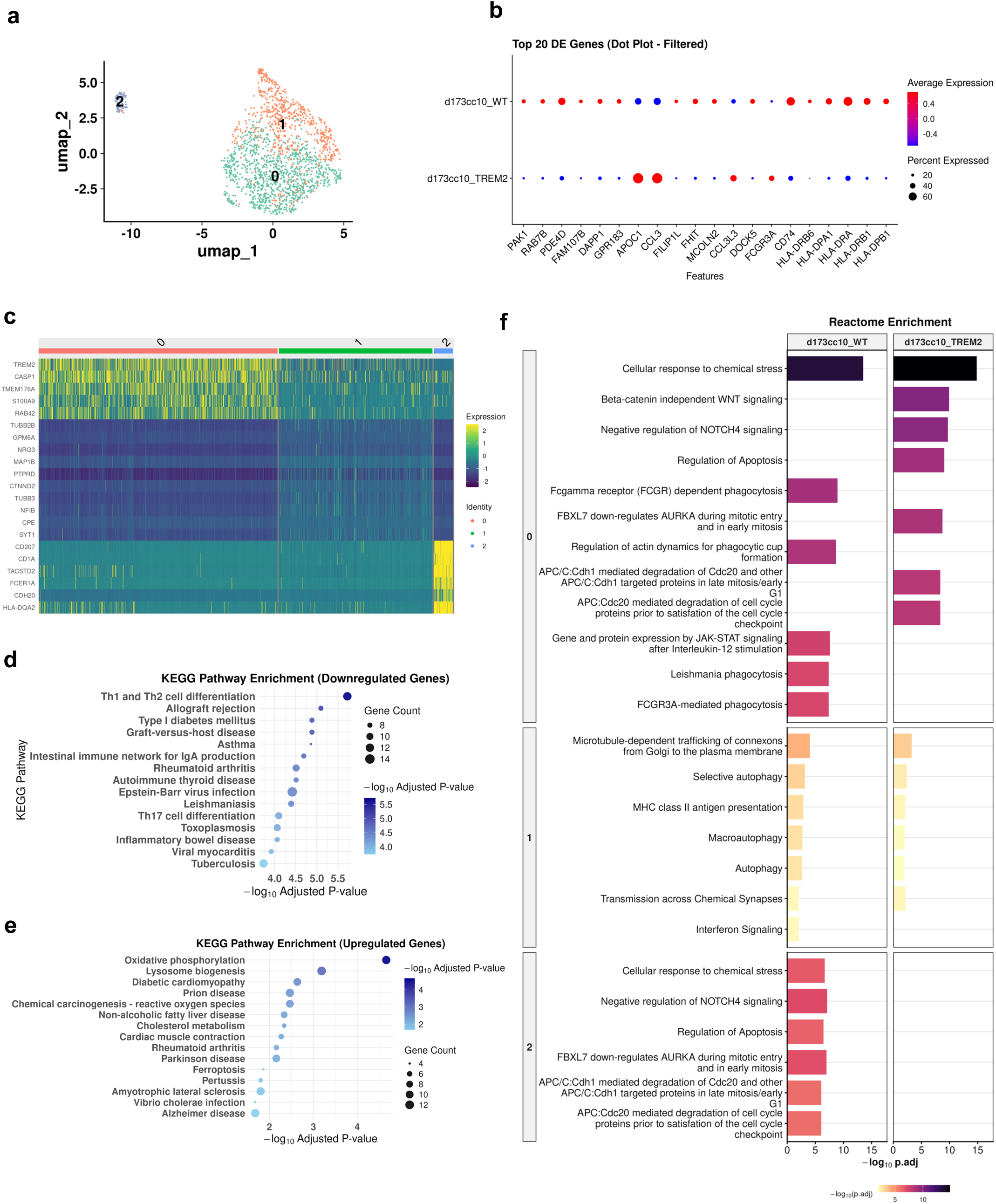
Microglia-specific subclustering and pathway alterations on day 173cc10. (a) UMAP visualization of microglial subclustering. (b) Dot plot of top 20 DEGs between *TREM2-R47H* and WT microglia. (c) Heatmap showing transcriptional differences between microglia subclusters (0, 1 and 2). (d-e) KEGG pathway enrichment analysis of differentially expressed genes in microglia comparing *TREM2-R47H* to WT organoids. (f) Comparative Reactome enrichment of cluster-specific marker genes. Bar plots illustrate the top enriched Reactome pathways for marker genes identified within each cluster, stratified by genotype (WT vs. *TREM2-R47H*). Genes were filtered for biological relevance to neurodegeneration, immune response, and metabolic state (Methods).

KEGG enrichment indicated suppression of gene modules related to T-cell differentiation, consistent with reduced antigen-presentation and adaptive immune interaction signatures. (Fig. 8d) (Supplementary Table 6). Upregulation, on the other hand, suggests stress-related and multiple neurodegenerative pathways including ferroptosis and AD-related enrichment (Fig. 8e). Consistent with KEGG, GO enrichment analyses of *TREM2-R47H* microglia (Supplementary Fig. 13 b-d) revealed marked suppression of MHC class II-mediated antigen processing and presentation, and adaptive immune response-related pathways including T cell differentiation, indicating reduced adaptive immune-associated programs in the mutant condition. The coordinated suppression of these pathways is consistent with the KEGG-based evidence of reduced antigen presentation capacity and impaired immune interaction in TREM2 mutant microglia.

Comparative Reactome analysis suggests that d173cc10 WT microglial clusters retain coordinated phagocytic, cytoskeletal, and cytokine-response programs, whereas *TREM2-R47H* microglia show reduced enrichment of these immune-clearance pathways and selective activation of alternative signaling modules such as β-catenin-independent WNT signaling. WT cultures also contained a distinct Cluster 2 enriched for stress-response, apoptosis-regulatory, and mitotic/cell-cycle control pathways, many of which were partially represented in Cluster 0, indicating that WT cells segregated into an additional specialized microglial state not resolved in *TREM2-R47H* cultures. This pattern is consistent with reduced microglial state diversity and altered functional organization in TREM2 mutant cells. All data have been reported in Supplementary table 12.

### Differential transcriptional responses to microglial co-culture in WT and TREM2-R47H organoids

To determine whether microglia induce genotype-specific effects on organoid cells, we compared transcriptional responses to microglial addition by evaluating DE between co-culture and microglia-negative conditions (d173cc10 vs d173) in WT and *TREM2-R47H* organoids, followed by interaction analysis (Fig. 9). Gene set enrichment analysis (GSEA) of interaction-ranked genes demonstrated that WT organoids preferentially activated mature neuronal and cortical programs following microglial co-culture, including neuronal subtype and neuro-supportive signatures (Fig. 9a). In contrast, *TREM2-R47H* organoids were enriched for mesenchymal-like and peripheral myeloid programs, indicating a divergent response characterized by partial loss of neural identity and acquisition of inflammatory-like states.

**Figure 9.**
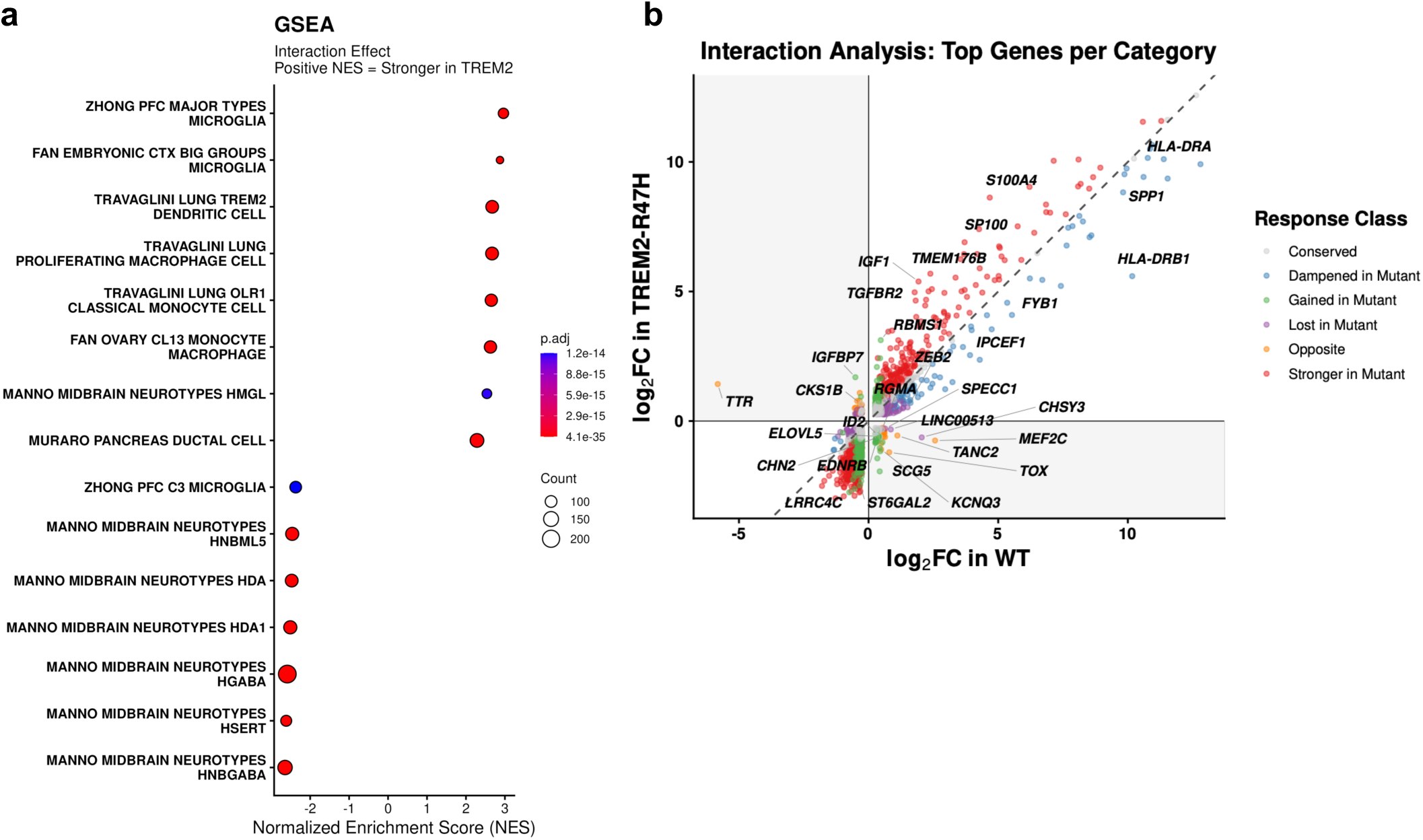
Interaction analysis comparing transcriptional responses to microglia addition in WT and TREM2-mutant organoids. (a) GSEA of interaction-ranked genes identifies enrichment of neuronal and cortical programs in WT, whereas TREM2 mutants show mesenchymal-like and peripheral myeloid signatures. (b) Scatter plot comparing log2 fold-change (d173cc10 vs d173) in WT (x-axis) and TREM2-mutant organoids (y-axis). Each point represents a gene. The diagonal line (y = x) indicates identical responses between genotypes. Gene-level comparison of log_2_ fold-changes reveals conserved, attenuated, and divergent responses, including genes lost, gained, or reversed in *TREM2-R47H*. Shaded quadrants highlight genes with discordant directional responses. Representative genes with the largest interaction effects (Δlog_2_ FC) are annotated.

Direct comparison of gene-level fold changes between WT and *TREM2-R47H* organoids further revealed both shared and genotype-specific responses to microglia (Fig. 9b). While a subset of genes responded similarly in both genotypes, *TREM2-R47H* organoids showed attenuated induction of antigen-presentation genes (*HLA-DRB1, HLA-DRA, FYB1*) and stronger upregulation of stress-associated genes (*S100A4, IGF1, TMEM176B*) following microglial co-culture. Additional responses were lost (*CHSY3, LINC00513, ZEB2*), gained (*RBMS1, SCG5, IGFBP7*), or directionally reversed, including increased *TTR* and decreased *MEF2C* and *TOX* relative to WT. Genes with the largest interaction effects implicated altered cellular identity, immune signaling, and stress-adaptive pathways. Together, these findings indicate that although microglia broadly influence both genotypes, TREM2 mutation markedly changes the quality and direction of specific transcriptional response.

### Development trajectory defined evident states of WT organoid cells over time and accumulation of uncommitted cells in TREM2 variant

To explore the dynamic progression of WT and TREM2 mutant samples, we performed pseudotime trajectory analysis using Monocle 2 based on single-cell transcriptomic profiles. First, we analyzed WT differentiation progression using samples from days 50, 100, 139, and 173 and cells from the earliest timepoint (50-day-old organoids) were assigned to the start of pseudotime. The analysis revealed a clear multibranch trajectory with three major cell states, suggesting divergent differentiation paths from progenitor-like cells toward either more mature neurons or glia (Fig. 10a-b).

**Figure 10.**
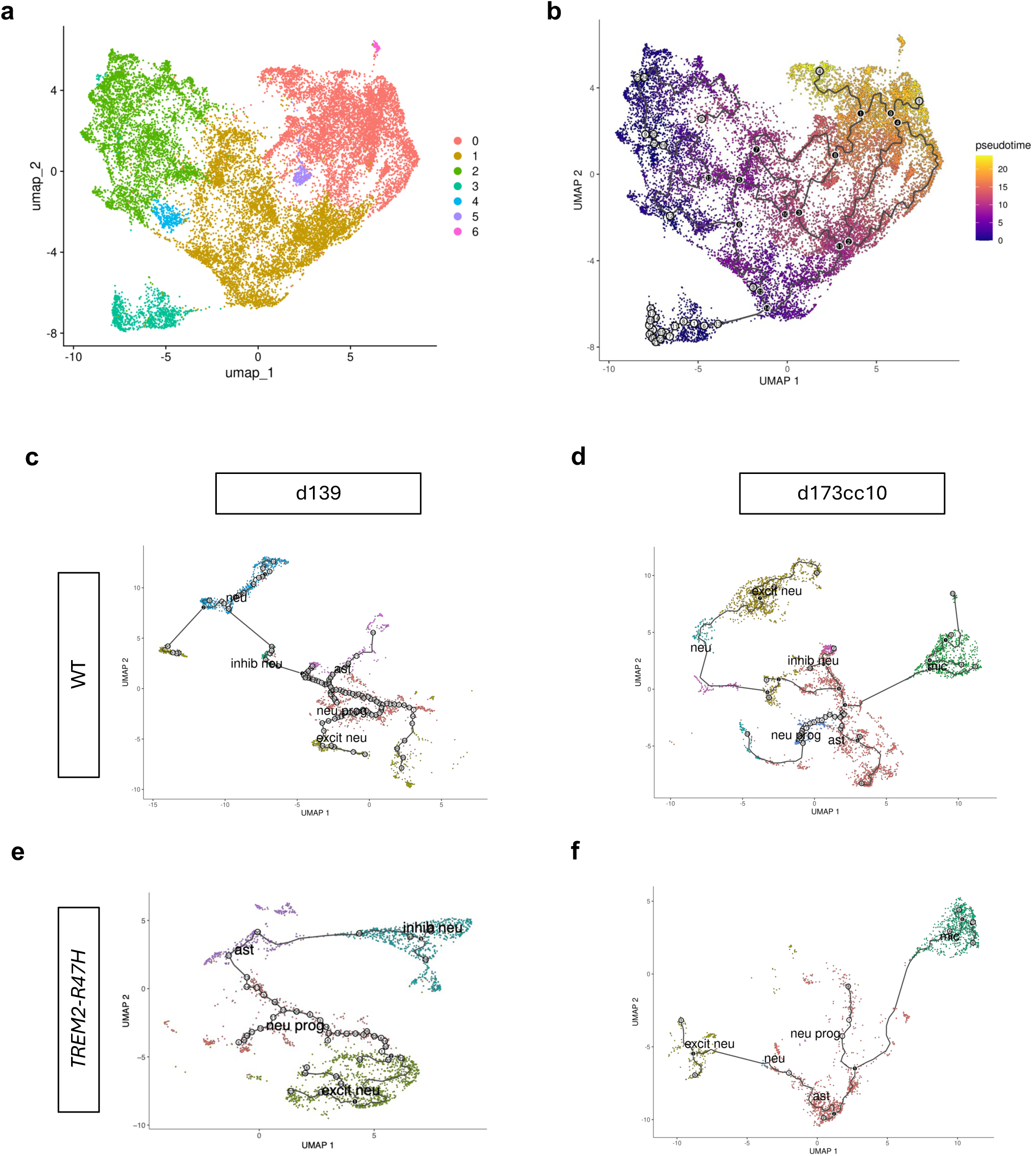
Pseudotime trajectory to analyze differentiation progression from scRNA-seq. (a-b) UMAP embedding of WT organoid cells integrated across developmental timepoints (days 50, 100, 139, and 173). (a) Cells colored by transcriptionally defined clusters. (b) Cells colored by pseudotime, illustrating the inferred differentiation trajectory and progressive maturation across time. UMAP projections illustrating the developmental or state-transition trajectories for WT (c, d) and *TREM2-R47H* (e, f) genotypes at days d139 and d173cc10, respectively.

To investigate the pseudotime trajectories between WT and *TREM2-R47H* organoids, d139 and microglia-containing (d173cc10) samples were analyzed. In both timepoints, *TREM2-R47H* samples exhibited altered or less coordinated cell fate commitment, whereas WT organoids displayed more distinct expression patterns with more ramified branching points (Fig. 10c-d).

Moreover, consistent with the loss of inhibitory neurons observed in our time-normalized subclustering, pseudotime trajectory analysis demonstrated a clear bifurcation failure in the *TREM2-R47H* background. While WT cells progressed along a branched trajectory toward mature inhibitory fates, the mutant path was truncated, with no cells on the inhibitory branch. Collectively, these results suggest that pseudotime analysis captures well-defined progression trajectories in WT organoids, while *TREM2-R47H* samples show disrupted transitions, highlighting key genes and pathways associated with these differences, though more analysis is needed to validate these results.

## Discussion

In this study, we employed a forebrain-microglia co-culture system to demonstrate that the *TREM2-R47H* mutation drives neurodegenerative environments across multiple cell populations. Our platform achieved homogeneous microglial internalization – contrasting with previous reports of peripheral-only integration^15^ – likely due to the use of individual U-bottom wells and the advanced maturation of the organoids at the time of co-culture. While the observed amoeboid microglial morphology suggests an activated or in vitro-adapted state^37^, the successful integration of these cells allowed for a high-resolution analysis of dynamic neuro-immune interactions.

A central finding is that *TREM2-R47H* influences neuronal and astrocytic states even in the absence of microglia, despite TREM2 expression being conventionally restricted to the myeloid lineage^39^. In monoculture, mutant organoids exhibited reduced neuronal proportions and transcriptional signatures of impaired maturation, shifting from synaptic assembly toward metabolic stress-response programs. Similarly, astrocytes in mutant organoids displayed reactive phenotypes or developmentally altered states. These "pre-microglia" alterations suggest that the TREM2 mutant background may influence early cellular programming or epigenetic landscapes, predisposing neural lineages to stress-adapted states.

The inhibitory neuron loss was observed upon time-normalized analysis. The initial presence of these clusters at day 139, followed by their depletion in later stages, mirrors the progressive nature of AD. The selective vulnerability of inhibitory neurons in our model is likely driven by a synergistic failure of transcriptional support. The downregulation of *MEF2C* and *RELN*, coupled with the loss of *ZEB2*-mediated lineage regulation, points to a fragile inhibitory circuit that is further compromised by the gain of stress-associated factors like *TTR* and *IGFBP7*. This molecular profile aligns with recent human AD brain studies identifying *MEF2C*^40,41^ and *RELN*^42^ as key nodes in the preservation of cognitive resilience, suggesting that the *TREM2-R47H* variant accelerates circuit collapse by undermining these protective networks.

The introduction of microglia further exacerbated these phenotypes. Immunohistochemical analysis revealed that while WT microglia successfully internalized pTau, mutant microglia exhibited impaired phagocytosis, confirming the effect of the R47H mutation in altering phagocytosis^43,44^. Notably, *TREM2-R47H* BOs exhibited a more extensive and spatially distributed pTau signal, which may reflect heightened inflammatory signaling associated with impaired microglial function, a process previously linked to tau hyperphosphorylation^45^.

Subclustering analysis identified a distinct, HLA-enriched microglial population in WT organoids – associated with antigen presentation and plaque clearance – that was absent in the *TREM2-R47H* model. The interaction analysis suggested that TREM2 mutation does not simply weaken the immune response but rewires it. While co-culture of WT organoids was associated with induction of antigen-presentation and homeostatic immune-response genes, *TREM2-R47H* organoids exhibited blunted immune signaling as well as disruption of normal neuronal function, shifting the tissue response away from supportive immune-neural interactions toward stress.

These results have significant translational implications. The early emergence of stress-associated signatures prior to microglial integration suggests that therapeutic windows for TREM2-related pathology may exist earlier in disease progression than previously thought. Furthermore, the ability of our co-culture platform to recapitulate these complex interactions makes it a tractable system for evaluating agonistic anti-TREM2 antibodies^46^ or other interventions aimed at restoring microglial homeostasis and study gene-specific mutations. However, these findings remain constrained by the neurodevelopmental nature of organoids and the potential for undetected CRISPR off-target effects in the absence of whole-genome sequencing.

In conclusion, we have leveraged a multi-timepoint single-cell approach to show that the *TREM2-R47H* variant exerts broad, cell-type-spanning effects that disrupt neuronal differentiation and astrocytic homeostasis. By demonstrating that this mutation affects more than just its canonical microglial target, our findings underscore the utility of complex organoid models in dissecting the interplay between cell-intrinsic and extrinsic drivers of AD. Future research integrating epigenomic profiling and longer-term co-cultures will be essential to determine how these early developmental shifts ultimately converge into the neurodegenerative phenotypes observed in the aging human brain.

## Methods

### Induced pluripotent stem cell (iPSC) maintenance

Two sets of induced pluripotent stem cells (iPSCs) from the WTC-11 cell line were obtained from Synthego (Redwood City, CA, USA). The first contained no genetic modifications and served as a WT control, while the other was purchased with a CRISPR-directed modification to contain the heterozygous *TREM2-R47H* mutation. CRISPR editing efficiency at the target locus was assessed by Sanger sequencing followed by ICE (Inference of CRISPR Edits) analysis (Synthego).

For iPSC culture, frozen WTC-11 iPSCs were thawed at 37°C and resuspended in mTeSR^TM^ Plus media (STEMCELL Technologies) supplemented with Y-27632 (ROCK inhibitor). iPSCs were plated on 60 mm cell culture plates coated with Matrigel® matrix (Corning) and cultured in an incubator with a humidified 37°C and 5% CO_2_. iPSCs were cultured as undifferentiated cells for 2 weeks with regular medium changes and passaged using Gentle Cell Dissociation Reagent (STEMCELL Technologies) upon reaching approximately 70% confluency.

### Generation and co-culture of brain organoids and microglia

iPSCs were used to form two sets of dorsal forebrain organoids, one with the *TREM2-R47H* mutant iPSCs and one with the WT iPSCs, using the STEMdiff™ Dorsal Forebrain Organoid Differentiation Kit (STEMCELL Technologies), following the manufacturer’s instructions. On day 0, iPSCs were seeded into AggreWell™800 plates (STEMCELL Technologies) at 4,000 cells per microwell. iPSCs were counted and concentrations were calculated using the Countess 3 Automated Cell Counter (Thermo Fisher Scientific). Plates were maintained in the same incubator environment as described in the iPSC maintenance section. On day 6, embryoid bodies were transferred to 6-well ultra-low adherent plates (STEMCELL Technologies) for suspension cultures with media change every 2 days. The organoid culture plates were kept on a shaker in the incubator and regularly shaken manually by lightly swirling the plates.

Throughout formation and maintenance, BOs were regularly observed under an inverted microscope and imaged to assess growth and viability. After day 43 of organoid formation, the BOs were maintained with regular medium changes as needed. Organoids were collected at days 50, 100, 139, 163 and 173 for analysis via single cell RNA sequencing, bulk RNA sequencing, immunofluorescence staining, and protein extractions, each of which is described below.

### Microglia generation

Human iPSCs, WTC-11 *TREM2-R47H* and WT control, were differentiated into microglia using the STEMdiff™ Hematopoietic Kit, followed by the STEMdiff™ Microglia Differentiation Kit and the STEMdiff™ Microglia Maturation Kit (STEMCELL Technologies), according to the manufacturer’s instructions. Briefly, iPSCs were first directed toward a hematopoietic progenitor fate using the STEMdiff™ Hematopoietic Kit. Resulting progenitors were subsequently differentiated into microglial precursors using the STEMdiff™ Microglia Differentiation Kit and further matured into microglia using the STEMdiff™ Microglia Maturation Kit. Cells were maintained under standard culture conditions as specified by the manufacturer.

### Organoid and microglia co-culture

The BOs and microglia were co-cultured by moving individual 163-day-old organoids into each well of a 96-well ultra-low adherent cell culture plate (Fisher Scientific), then introducing 28-day-old microglia into the wells at a concentration of 83,000-90,000 microglia per organoid. A partial media change was performed every two days, with care taken not to disturb the microglia cells at the bottom of the plate. The co-culture was imaged daily to track microglia integration into the organoids. On the tenth day of the co-culture, when BOs were 173 days old, all BOs were collected for analysis.

### Cryosectioning and immunofluorescence staining

For cryosectioning, BOs were collected from culture and rinsed three times in phosphate-buffered saline (PBS), then fixed overnight at 4°C in 4% paraformaldehyde (PFA) in PBS. Following fixation, BOs were cryoprotected by immersion in 30% sucrose in PBS overnight at 4°C, embedded in OCT compound containing 30% sucrose, and stored at -80°C. BOs were cryosectioned at a thickness of 10 µm, and sections were mounted onto positively charged microscope slides. Sections were stored at -80°C until staining.

For immunofluorescence staining, mounted organoid sections were equilibrated to room temperature and rinsed in PBS containing 0.1% Tween-20 (PBST). Sections were post-fixed in 4% PFA/PBS for 10 minutes at room temperature, then washed in PBST. Sections were blocked in 5% bovine serum albumin (BSA) in PBS for 1 hour at room temperature, followed by incubation with primary antibodies diluted in blocking buffer in a humidified chamber at 4°C overnight. The following primary antibodies were used: anti-βIII-tubulin (clone TUJ1) (STEMCELL Technologies, cat# 60052, 1:500), Phospho-Tau (Ser202, Thr205) Monoclonal Antibody (AT8) (ThermoFisher Scientific, cat# MN1020, 1:100), Iba-1 Polyclonal Antibody (Invitrogen, cat# PIPA527436, 1:200), anti-GFAP (clone 2E1.E9) (STEMCELL Technologies, cat# 60048.1, 1:100), Aβ (MOAB-2) (Novus Biologicals, cat# NBP2-13075, 1:200). After primary antibody incubation, sections were washed and incubated with species-appropriate fluorescent secondary antibodies diluted in blocking buffer for 2 hours at room temperature in the dark. Cell nuclei were counterstained with VECTASHIELD PLUS Antifade Mounting Medium with 4′,6-diamidino-2-phenylindole (DAPI) mounting medium (Vector Laboratories, cat# H-2000-10).

Stained sections were stored at 4°C in the dark and imaged using a Zeiss LSM confocal microscope with Airyscan super-resolution detection (Carl Zeiss Microscopy).

### Bulk RNA extraction and sequencing

Total RNA was extracted from BOs using the Maxwell® RSC simplyRNA Cells/Tissue Kit (Promega) following the manufacturer’s instructions. Briefly, organoids were collected from culture medium, centrifuged to remove residual medium, and homogenized in 200 µL of cold 1-Thioglycerol/Homogenization Solution (Promega). 200 µL of lysis buffer was added prior to loading samples onto the Maxwell® RSC instrument for automated RNA extraction.

Extracted RNA was quantified using a Qubit fluorometer (Thermo Fisher Scientific), and 100 ng of total RNA per sample was used for library preparation with the KAPA RNA HyperPrep Kit with RiboErase (HMR) (Roche). RNA and library quality were assessed using the TapeStation 4200 (Agilent Technologies). Libraries were sequenced on an Illumina NovaSeq X Plus platform to a depth of approximately 50 million paired-end reads per sample.

### scRNA - 10X genomics

A custom dissociation buffer was prepared in Hank’s Balanced Salt Solution (HBSS) containing 15 U/mL activated papain and 60 U/mL DNase I (STEMCELL Technologies). Papain activation buffer consisted of 1.1 mM EDTA, 67 µM β-mercaptoethanol, and 5.5 mM L-cysteine HCl. An ovomucoid protease inhibitor solution was prepared by dissolving 10 mg ovomucoid protease inhibitor in 1 mL HBSS.

Organoids were collected from culture, and the medium was removed. Organoids were incubated in 500 µL dissociation buffer on a shaker at 250 rpm and 37°C for 30 min, followed by gentle trituration and an additional 10-minute incubation at 300 rpm and 37°C. Samples were briefly pipetted to further dissociate remaining tissue fragments, and 1 mL of the cell suspension was immediately transferred into a tube containing 1 mL ovomucoid protease inhibitor solution to quench enzymatic activity.

Samples were centrifuged at 500 × g for 5 min at 4°C. Supernatants were carefully removed without disturbing the pellet, and cells were resuspended in 1 mL HBSS and filtered through a Flowmi™ 40 µm cell strainer (pluriSelect-USA Inc.).

Cell viability and concentration were assessed by mixing 10 µL of cell suspension with 10 µL of 0.4% Trypan Blue (Invitrogen) and loading the mixture onto chamber slides in duplicate. Cell counts were obtained using a Countess™ 3 FL Automated Cell Counter (Invitrogen). 6,000 viable cells per sample were loaded to generate gel bead–in-emulsions (GEMs) using the Chromium X Controller with Single Cell 3′ Reagent Kits v3.1 Dual Index (10x Genomics), following the manufacturer’s instructions.

Library quality was assessed using a TapeStation 4200 (Agilent Technologies). Library rebalancing was performed based on iSeq read distributions per sample, and pooled libraries were sequenced on an Illumina NovaSeq X Plus platform to a depth of approximately 20,000 reads per cell.

### Data analysis

#### scRNA-seq data preprocessing, clustering, and cell type annotation

Gene expression matrices were generated by aligning sequencing reads to the 10x Genomics GRCh38 reference genome (refdata-gex-GRCh38-2024-A) using Cell Ranger v8.0 (10x Genomics) with default settings. Ambient RNA contamination was removed using CellBender (v0.3.0) with default parameters.

The resulting denoised matrices were imported into Seurat (v5.3) for downstream analysis. Individual samples were merged into a single object, and initial filtering was performed to remove low-quality features and droplets. Specifically, genes expressed in fewer than 3 cells and cells expressing fewer than 200 unique genes were excluded. Further QC filtering was applied to retain only high-quality cells with: 200 < nFeature_RNA < 6,000; 500 < nCount_RNA < 25,000; and a mitochondrial content < 15%.

Data were processed and variance-stabilized within the SCTransform framework in Seurat. To preserve stage-specific biological signals and account for technical variation, a tiered normalization strategy was employed: d139 samples were normalized independently as a single cohort, while d173 and d173cc10 samples were grouped and normalized together to provide a shared mathematical space for microglia integration analysis. Mitochondrial gene expression (percent.mt) was regressed out during the normalization process to minimize technical noise.

For the global integration and clustering of all timepoints, dimensionality reduction via UMAP, nearest-neighbor graph construction, and Louvain clustering were performed using the first 50 principal components (PCs) with a clustering resolution of 0.1.

Cell type identities were assigned manually at the cluster level using canonical marker genes reported in published single-cell/single-nucleus datasets. Cluster-specific marker genes were identified using Seurat’s FindAllMarkers (positive markers only; min.pct = 0.25, logfc.threshold = 0.25) on the SCTransform assay. Clusters were annotated by examining the expression of curated markers for astrocytes (*AQP4, GFAP*), neurons (*SYT1, SNAP25*), excitatory neurons (*SATB2, SLC17A7*), inhibitory neurons (*GAD1, GAD2*), microglia (*ADAM28, APBB1IP*), and neural progenitor cells (*SOX2, PAX6*).

To evaluate the transcriptomic impact of the mutation, pairwise WT and *TREM2-R47H* samples from the same timepoint were subsetted for DE analysis. Single-cell RNA-seq samples consisted of pooled material from 3–5 organoids per condition. Because independent sequencing replicates were not available for each condition, DE analyses were performed at the single-cell level using Seurat FindMarkers with the Wilcoxon rank-sum test. Genes detected in at least 10% of cells were retained for testing, and a minimum log2 fold-change threshold of 0.25 was applied. P-values were adjusted for multiple comparisons using Bonferroni correction, and genes with adjusted p < 0.05 were considered significant. Results were interpreted as cell-state-associated transcriptional differences and supported by pathway-level analyses. This batch-aware normalization and subsetting approach allowed for high-sensitivity detection of cell-intrinsic defects at day 139 and genotype-specific microglia effect at day 173/cc10. Subsequent functional profiling, including pathway enrichment (GO, KEGG, and Reactome) and interaction analysis, were performed within each paired dataset to characterize the functional divergence between genotypes.

To resolve fine-grained functional states and subtle genotype-specific transcriptomic shifts, lineage-specific subclustering was performed on major cell populations (microglia, neurons, neural progenitor cells and astrocytes). For each lineage, relevant cells were subsetted from the global integrated object and subjected to an independent round of analysis to minimize ’background’ variance from unrelated cell types. This included recalculated variable feature selection, data scaling, and PCA. High-resolution UMAP projections and Louvain clustering were then performed using a lineage-optimized number of PCs (top 20) and 0.1 resolution parameters. These high-resolution subclusters provided the basis for all downstream cluster-specific marker identification, Reactome pathway enrichment, and genotype-interaction analyses. For main figure visualizations, Reactome results were filtered using a keyword-based strategy to prioritize pathways related to immune signaling and neurodegeneration (keywords included: "immune", "inflamm", "cytokine", "chemokine", "Toll-like", "phagocytosis", "interferon", "complement", "antigen presentation", "innate immunity", "adaptive immunity", "mic", "neurodegeneration", "Alzheimer", "tau", "amyloid", "oxidative", "apoptosis", "stress", "lysosome", "autophagy", "clearance", "cell migration", "response to wounding", "signal transduction", "MAPK", "JAK-STAT", "NF-kB", "Notch", "PI3K", "TGF-beta", "Wnt", "cell cycle", "proliferation", "DNA replication", "mitosis", "glycolysis", "oxidative phosphorylation", "mitochondrial", "fatty acid metabolism", "cholesterol", "lipid") (Full unfiltered results provided in Supplementary Table X).

### Genotype-specific interaction analysis of microglial responses

To quantify genotype-dependent differences in the transcriptional response to microglia, differential expression contrasts generated as described above were performed separately in WT and *TREM2-R47H* organoids (microglia-positive vs microglia-negative within genotype). Gene-level log_2_ fold-changes and adjusted p-values from each contrast were merged by gene symbol, and an interaction score was calculated as:

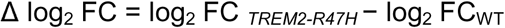

where positive values indicate a stronger transcriptional induction in *TREM2-R47H* relative to WT, and negative values indicate a stronger response in WT.

Genes were further classified into discrete response categories according to statistical significance in each genotype, concordance of response direction, and relative effect magnitude. These categories included conserved responses, enhanced responses in mutant cells, attenuated responses in mutant cells, genotype-specific gains or losses of responsiveness, and directionally reversed responses.

To visualize gene-level concordance between genotypes, WT and *TREM2-R47H* log_2_ fold-changes were plotted against one another in a two-dimensional interaction scatter plot. The identity line (y = x) denotes equivalent responses across genotypes, whereas deviation from this line reflects genotype-dependent modulation of the microglial response. Genes with the largest absolute interaction scores were annotated.

To identify biological programs differentially modulated by genotype during microglial co-culture, genes were rank-ordered by the interaction score (Δlog_2_FC) and subjected to preranked GSEA using clusterProfiler. Human MSigDB gene sets were retrieved using msigdbr. Gene sets containing 10-500 genes were tested, and significance was assessed using Benjamini-Hochberg adjusted p-values (FDR < 0.05). Positive normalized enrichment scores (NES) indicate pathways more strongly induced in *TREM2-R47H* organoids, whereas negative indicate pathways preferentially induced in WT organoids.

### Pseudotime analysis

Pseudotime analysis was performed to examine transcriptional progression across developmental time points and between genotypes. Cells from scRNA-seq datasets were ordered along inferred trajectories using Monocle 2, which reconstructs trajectories based on gene expression dynamics. Pseudotime values were calculated for individual cells and overlaid onto UMAP embeddings generated in Seurat for visualization.

For developmental progression analysis, WT organoid samples collected at days 50, 100, 139, and 173 were included. For genotype comparison, WT and *TREM2-R47H* co-cultured organoid samples collected on the same day were analyzed. Pseudotime values were used solely for visualization of transcriptional progression and comparison of inferred trajectories between conditions.

### Nextflow for bulk RNA-seq

Bulk RNA-seq data were processed using the nf-core/rnaseq pipeline implemented in Nextflow (version 24.10.3). The pipeline was executed using Singularity containers. Reads were aligned to the human reference genome GRCh38 using the primary assembly FASTA file, and gene-level quantification was performed using GENCODE annotation release 112. Default pipeline parameters were used unless otherwise specified.

### Bulk RNA-seq downstream analysis

Gene-level count matrices generated by the nf-core/rnaseq pipeline were imported into R (version 3.14.0) for downstream analysis. Sample metadata were curated to include genotype (WT or TREM2* i.e. *TREM2-R47H*) and developmental time point (days 100, 163, and 173).

For exploratory analysis, variance stabilizing transformation (VST) was performed using DESeq2, and principal component analysis (PCA) was conducted on VST-transformed counts to assess global transcriptional differences across genotypes and time points.

Differential gene expression analysis was performed using DESeq2 with pairwise comparisons between WT and *TREM2-R47H* samples conducted separately at each time point for days 100, 163, and 173 (cc10) (DEGs pval.adj < 0.05, log2(FC) = 0.25, min.pct = 0.1, test = wilcox). Lowly expressed genes were filtered prior to analysis by retaining genes with a mean count ≥10 across samples. Differential expression testing was performed using the DESeq2 Wald test, and statistical significance was assessed using nominal P values. The specific comparisons performed are indicated in the figure legends.

Stress-related transcription was assessed by quantifying bulk RNA-seq expression of a predefined panel of 10 genes associated with hypoxia, metabolic stress, ER stress, and apoptotic signaling (*HIF1A, BAX, BNIP3, CASP3, LDHA, HSPA5, VEGFA, DDIT3, SLC2A1, ATF4*) across four developmental time points starting at day 100, with an aggregate stress score computed as the mean expression of this gene set per sample.

### Statistical analysis

Statistical analyses were performed using R (version 3.14.0). Unless otherwise stated, data are presented as mean ± standard error of the mean (SEM). Statistical significance was assessed using two-sided tests. P values < 0.05 were considered statistically significant.

### AI-assisted writing and editing

During the preparation of this manuscript, the authors used AI-assisted technology to perform spelling and grammar checks and to refine the prose for better readability. The tool was further employed to ensure the technical narrative was accessible to a broad, multidisciplinary scientific audience. Following this process, the authors independently reviewed, edited, and verified all output to ensure accuracy. The authors take full responsibility for the final content of the manuscript.

## Data availability

The raw and processed scRNA-seq and bulk RNA-seq datasets have been deposited in the Gene Expression Omnibus (GEO) and will be publicly available after publication. All other data supporting the findings of this study are included within the article and its Supplementary Information files.

## Code availability

The code for processing the data, performing the analyses, generating figures, and reproducing the results presented in this study is available at GitHub (link will be released after publication).

## Acknowledgements

Support from the Arizona Alzheimer’s Consortium and Arizona’s Department of Health Services in part funded this work. The authors thank Dr Ignazio Piras for his comments on this manuscript.

## Author contributions

MJH supervised the study and applied for funding. ASK, MJH designed the study and experiments. ASK, KEL, AA performed experiments. ASK, FE, MJH analyzed and interpreted data. ASK, KEL, FE wrote the manuscript. ASK, KEL, FE, MJH edited the manuscript. All authors read and approved the final version of the manuscript.

## Competing interests

The authors declare no competing interests.

## Notes

### Competing Interest Statement

The authors have declared no competing interest.

### Summary of Updates

This revised version includes updated analyses, figure refinements, and textual revisions made during manuscript preparation for journal submission. Several datasets were reprocessed using an improved analytical pipeline, resulting in minor updates to quantitative results and visualizations. These changes do not alter the overall conclusions of the study but strengthen the robustness and clarity of the findings.

