## Supplementary Figures for "The TREM2-R47H Variant Drives Alzheimer’s-Relevant Alterations in Forebrain Organoids Beyond Microglial Populations"

### Slide 1
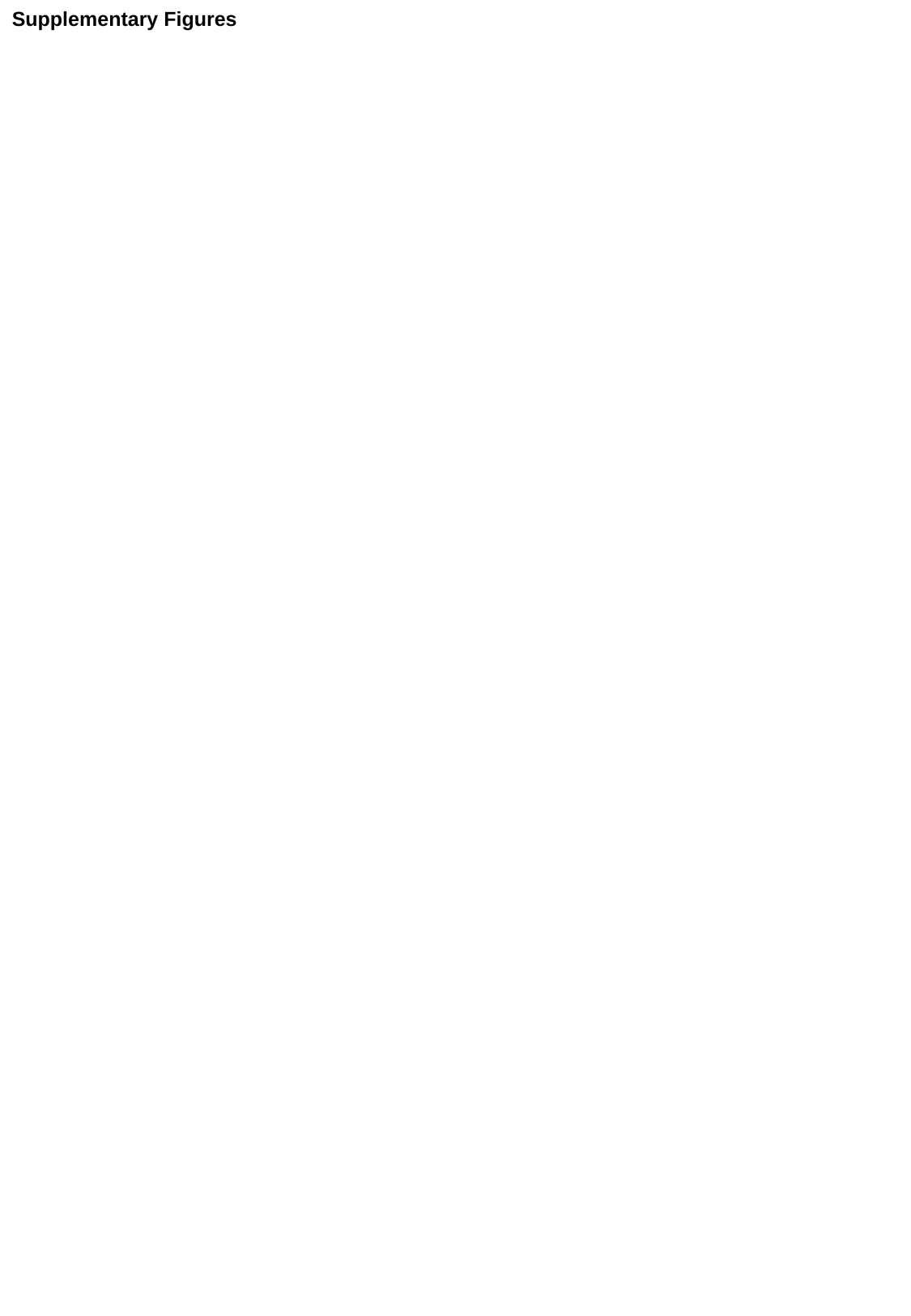

Supplementary Figures

### Slide 2
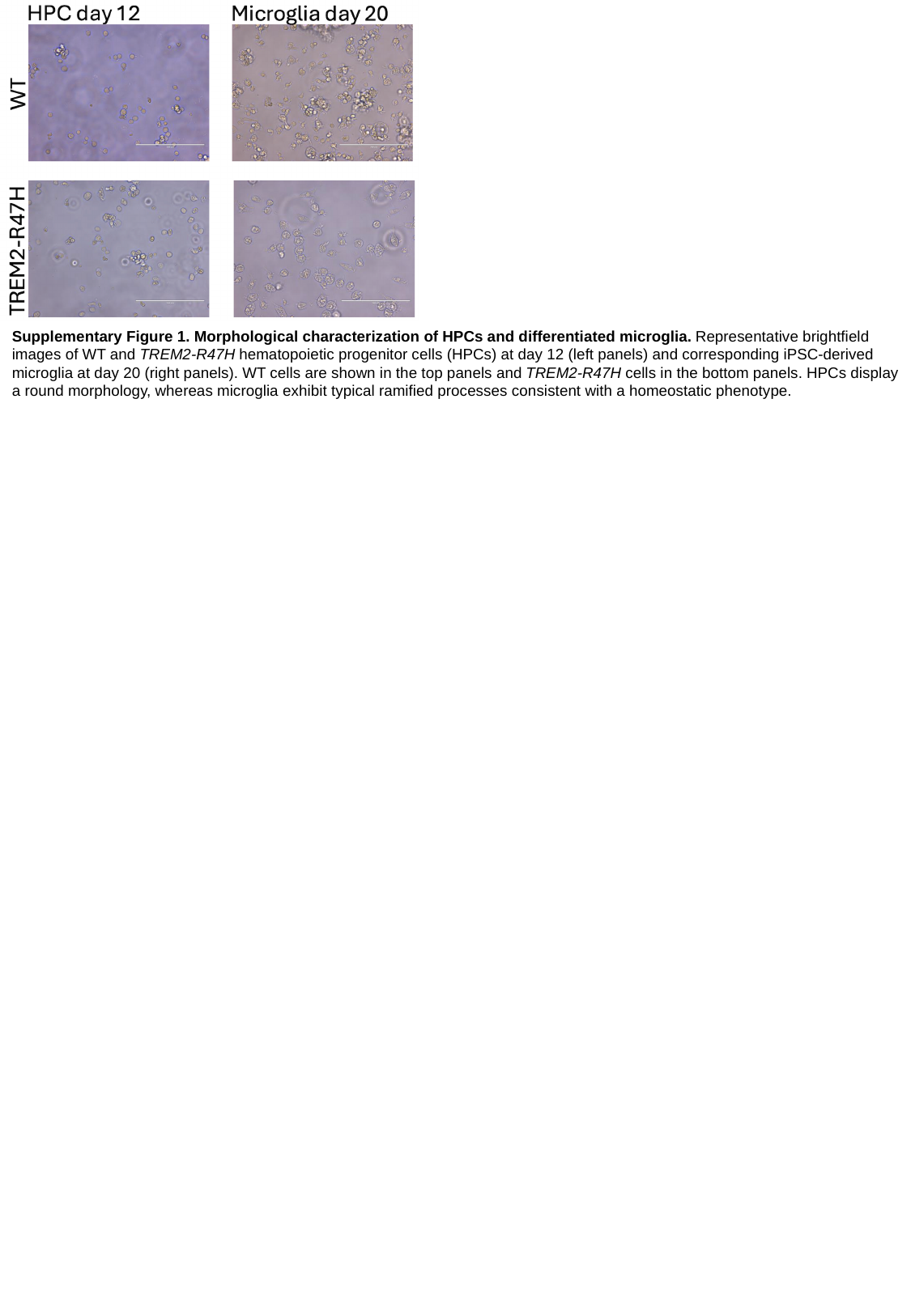

Supplementary Figure 1. Morphological characterization of HPCs and differentiated microglia. Representative brightfield images of WT and TREM2-R47H hematopoietic progenitor cells (HPCs) at day 12 (left panels) and corresponding iPSC-derived microglia at day 20 (right panels). WT cells are shown in the top panels and TREM2-R47H cells in the bottom panels. HPCs display a round morphology, whereas microglia exhibit typical ramified processes consistent with a homeostatic phenotype.

### Slide 3
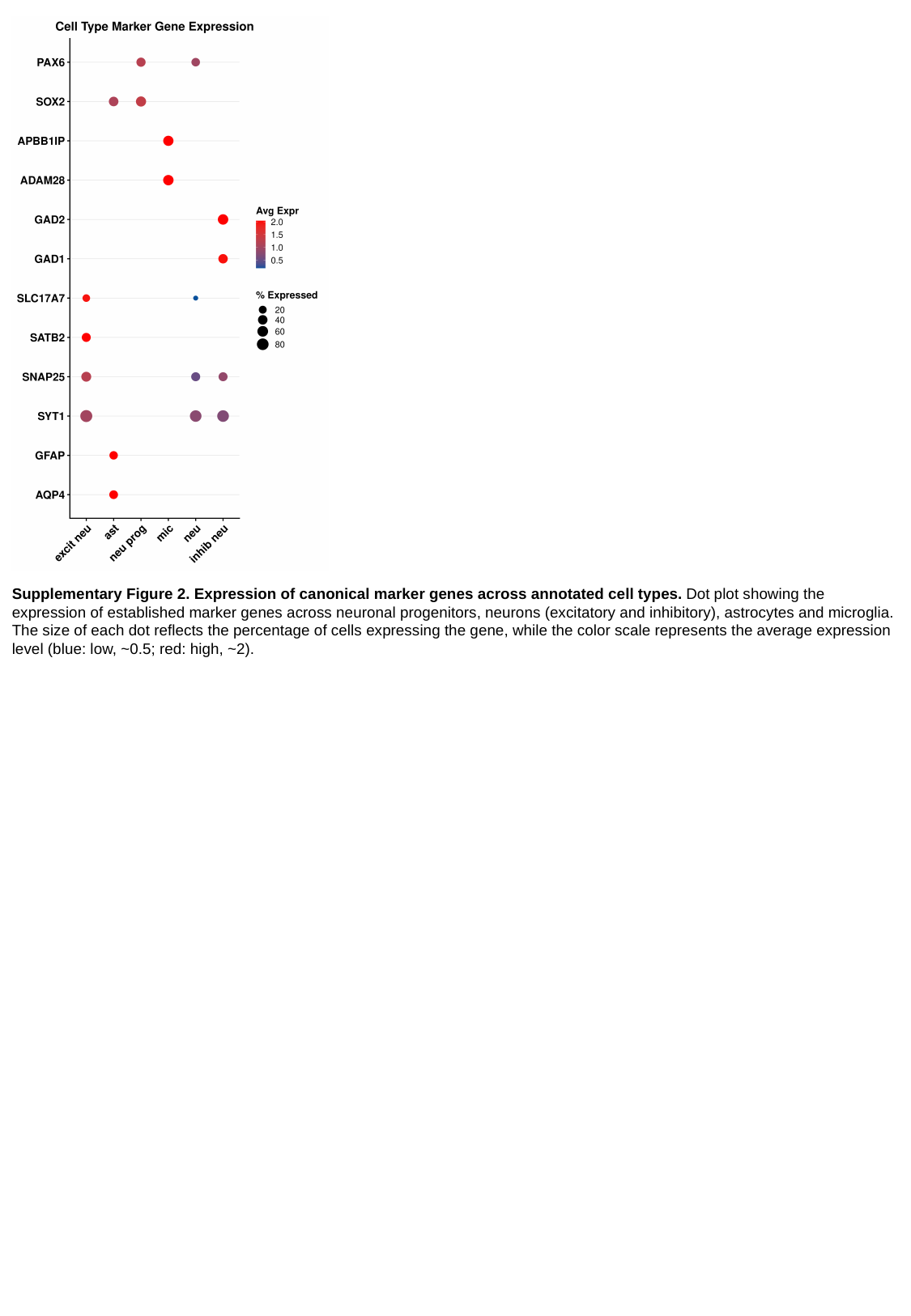

Supplementary Figure 2. Expression of canonical marker genes across annotated cell types. Dot plot showing the expression of established marker genes across neuronal progenitors, neurons (excitatory and inhibitory), astrocytes and microglia. The size of each dot reflects the percentage of cells expressing the gene, while the color scale represents the average expression level (blue: low, ~0.5; red: high, ~2).

### Slide 4
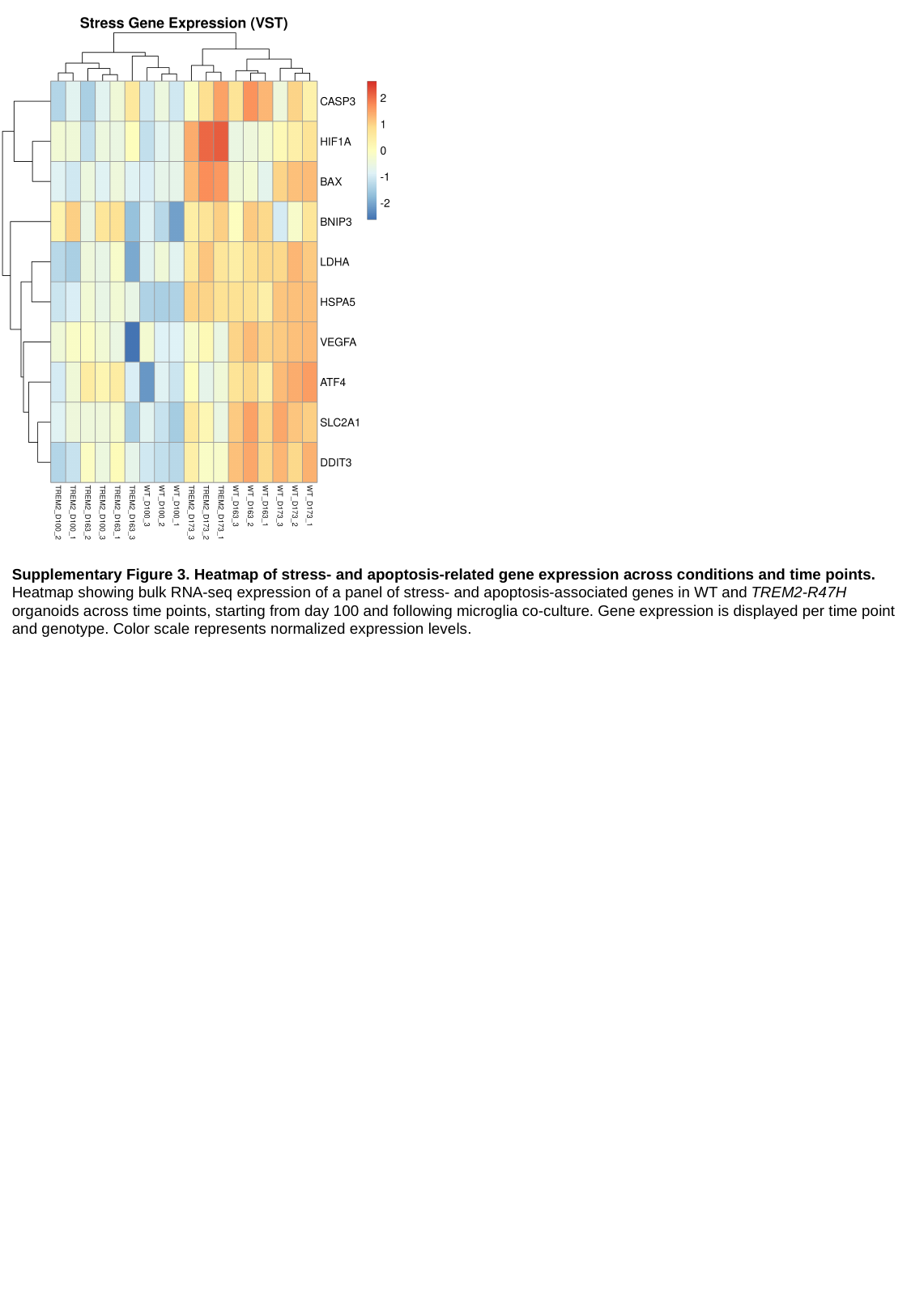

Supplementary Figure 3. Heatmap of stress- and apoptosis-related gene expression across conditions and time points. Heatmap showing bulk RNA-seq expression of a panel of stress- and apoptosis-associated genes in WT and TREM2-R47H organoids across time points, starting from day 100 and following microglia co-culture. Gene expression is displayed per time point and genotype. Color scale represents normalized expression levels.

### Slide 5
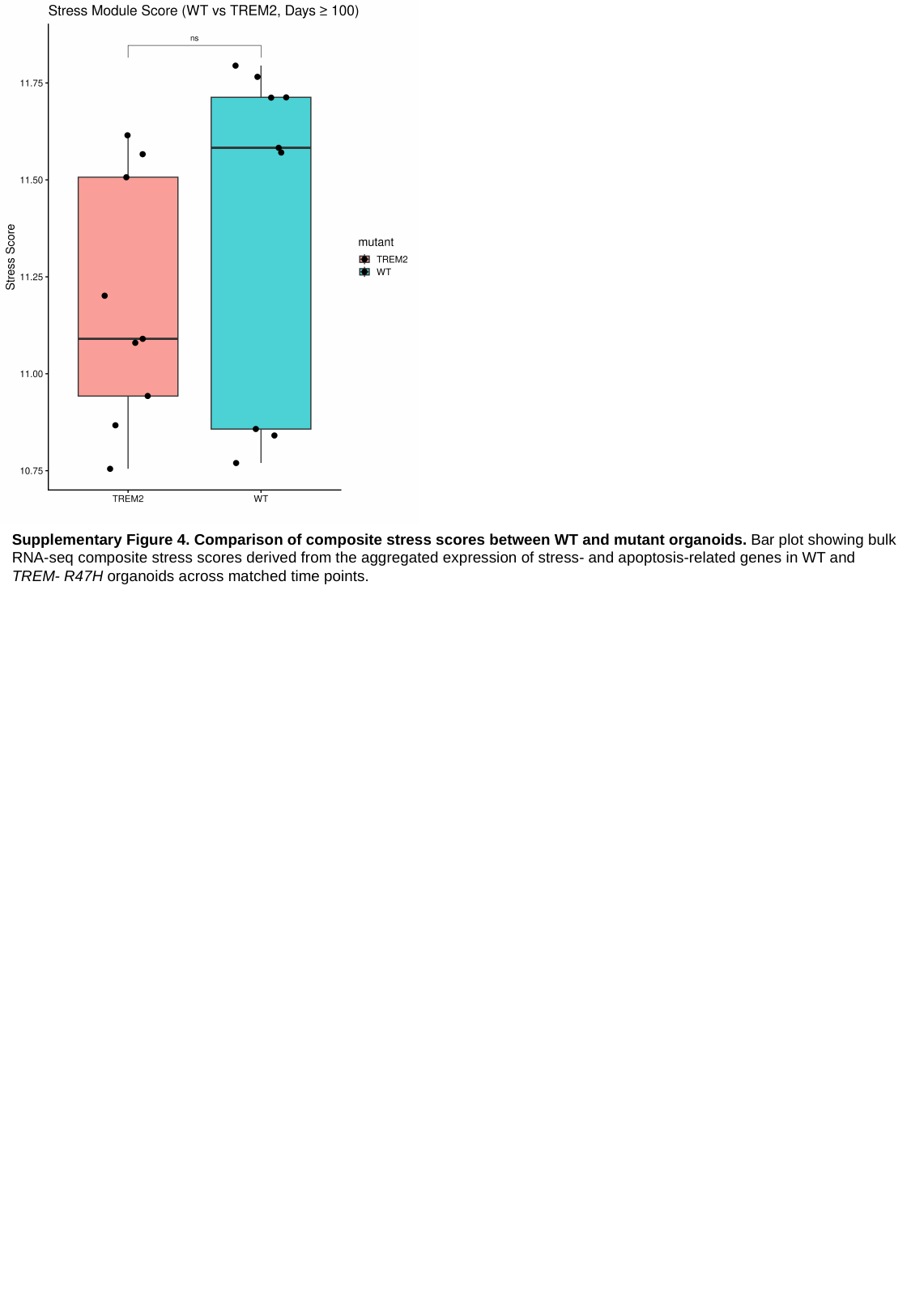

Supplementary Figure 4. Comparison of composite stress scores between WT and mutant organoids. Bar plot showing bulk RNA-seq composite stress scores derived from the aggregated expression of stress- and apoptosis-related genes in WT and TREM- R47H organoids across matched time points.

### Slide 6
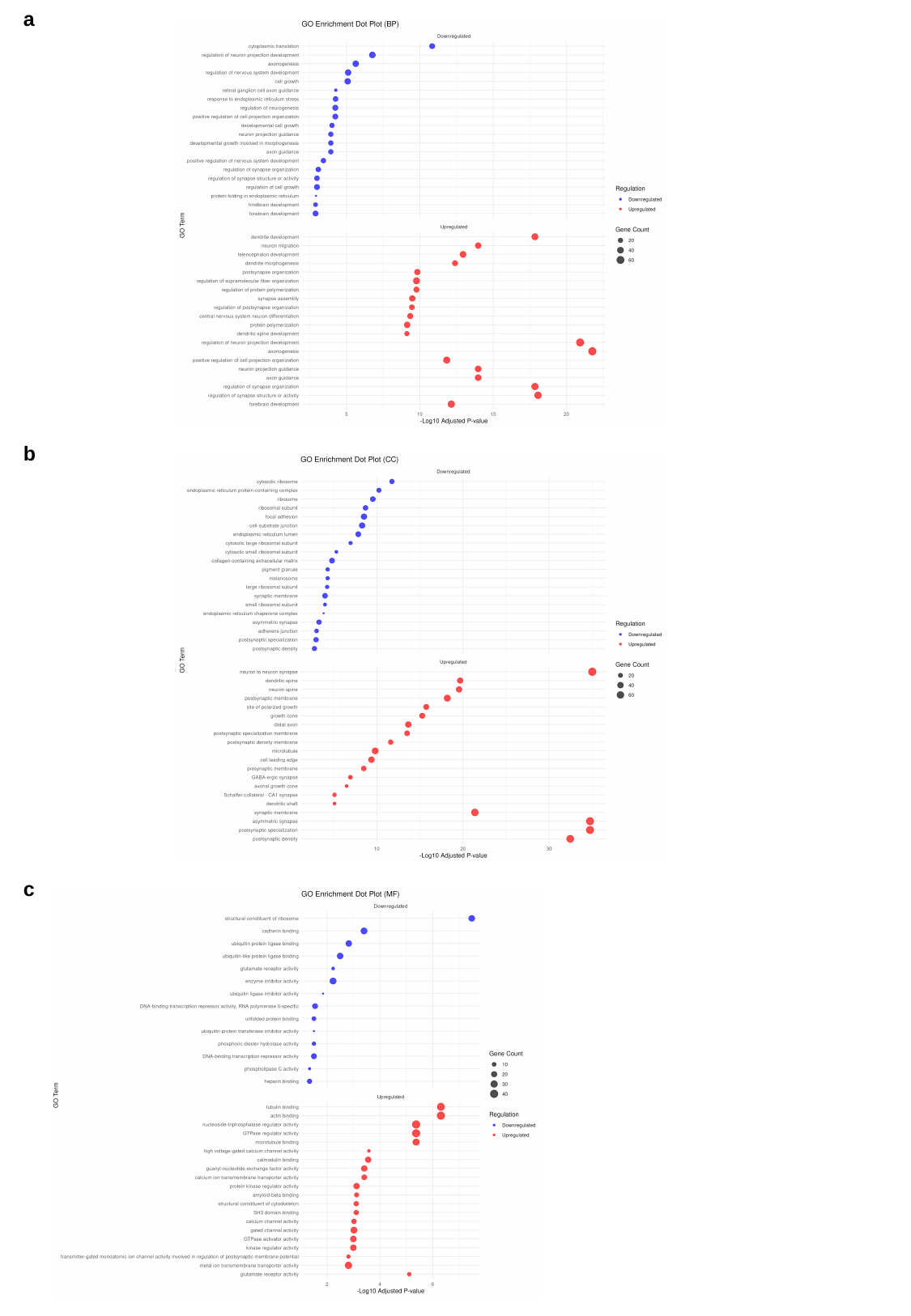

a
b
c

### Slide 7
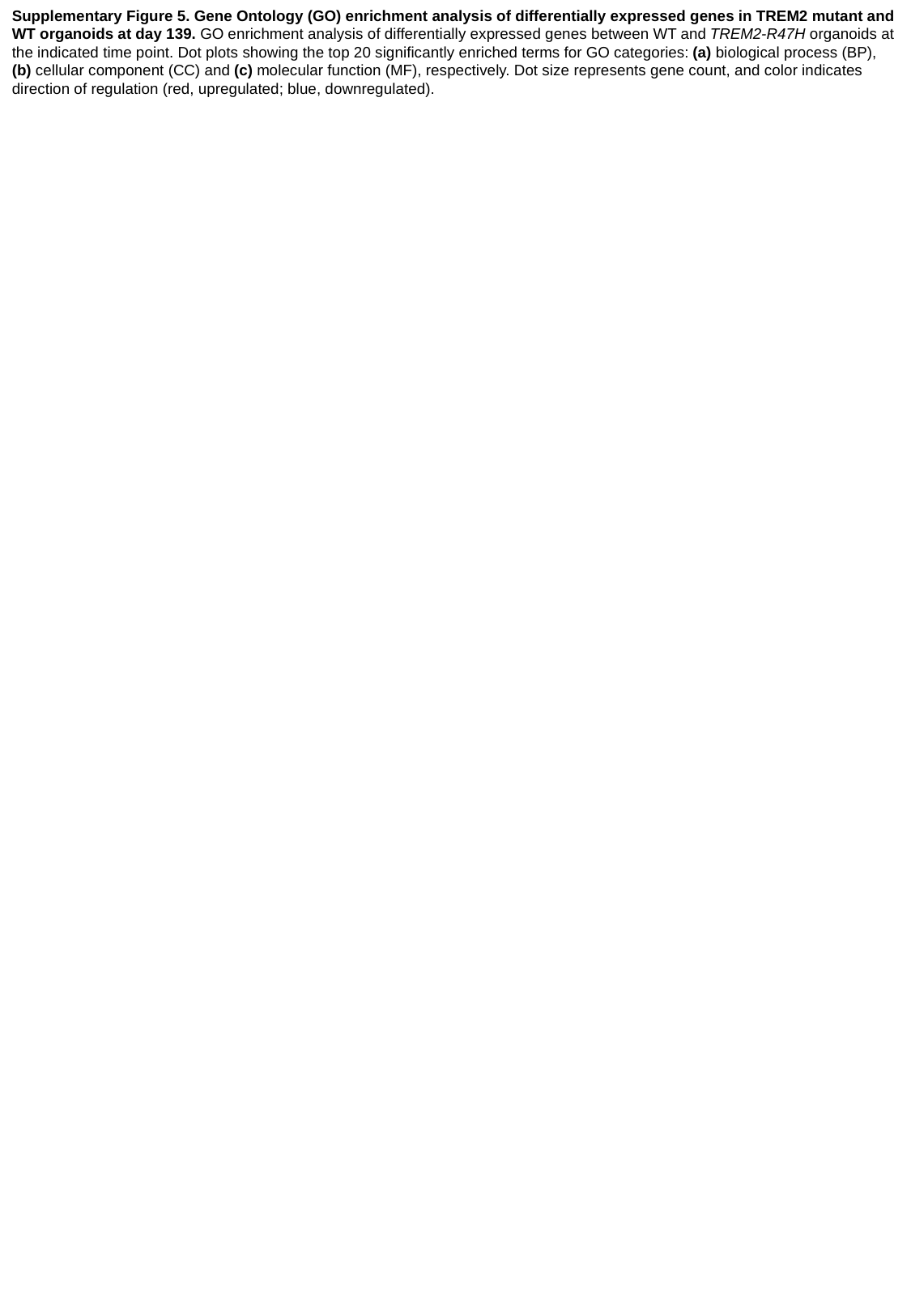

Supplementary Figure 5. Gene Ontology (GO) enrichment analysis of differentially expressed genes in TREM2 mutant and WT organoids at day 139. GO enrichment analysis of differentially expressed genes between WT and TREM2-R47H organoids at the indicated time point. Dot plots showing the top 20 significantly enriched terms for GO categories: (a) biological process (BP), (b) cellular component (CC) and (c) molecular function (MF), respectively. Dot size represents gene count, and color indicates direction of regulation (red, upregulated; blue, downregulated).

### Slide 8
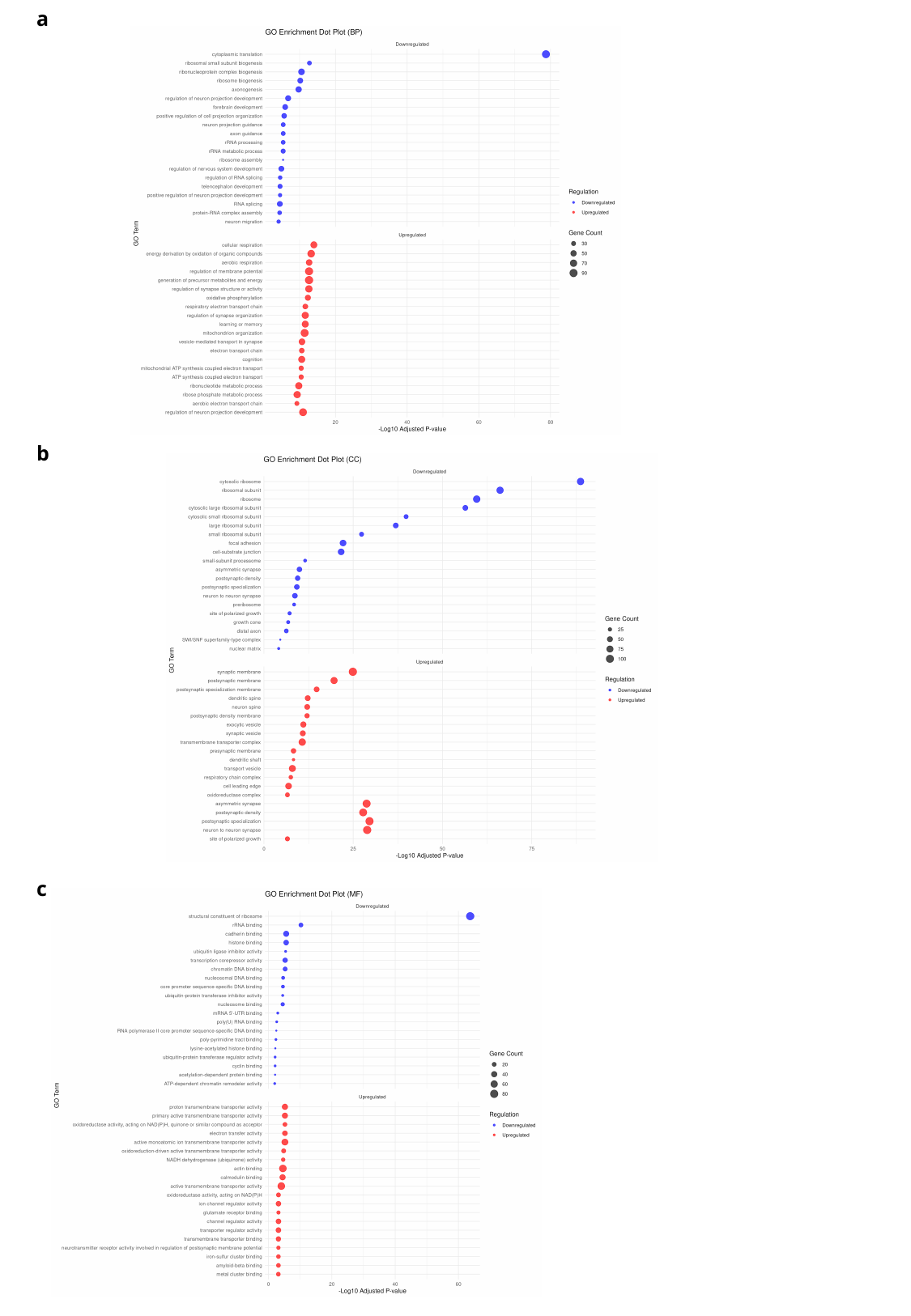

a
b
c

### Slide 9
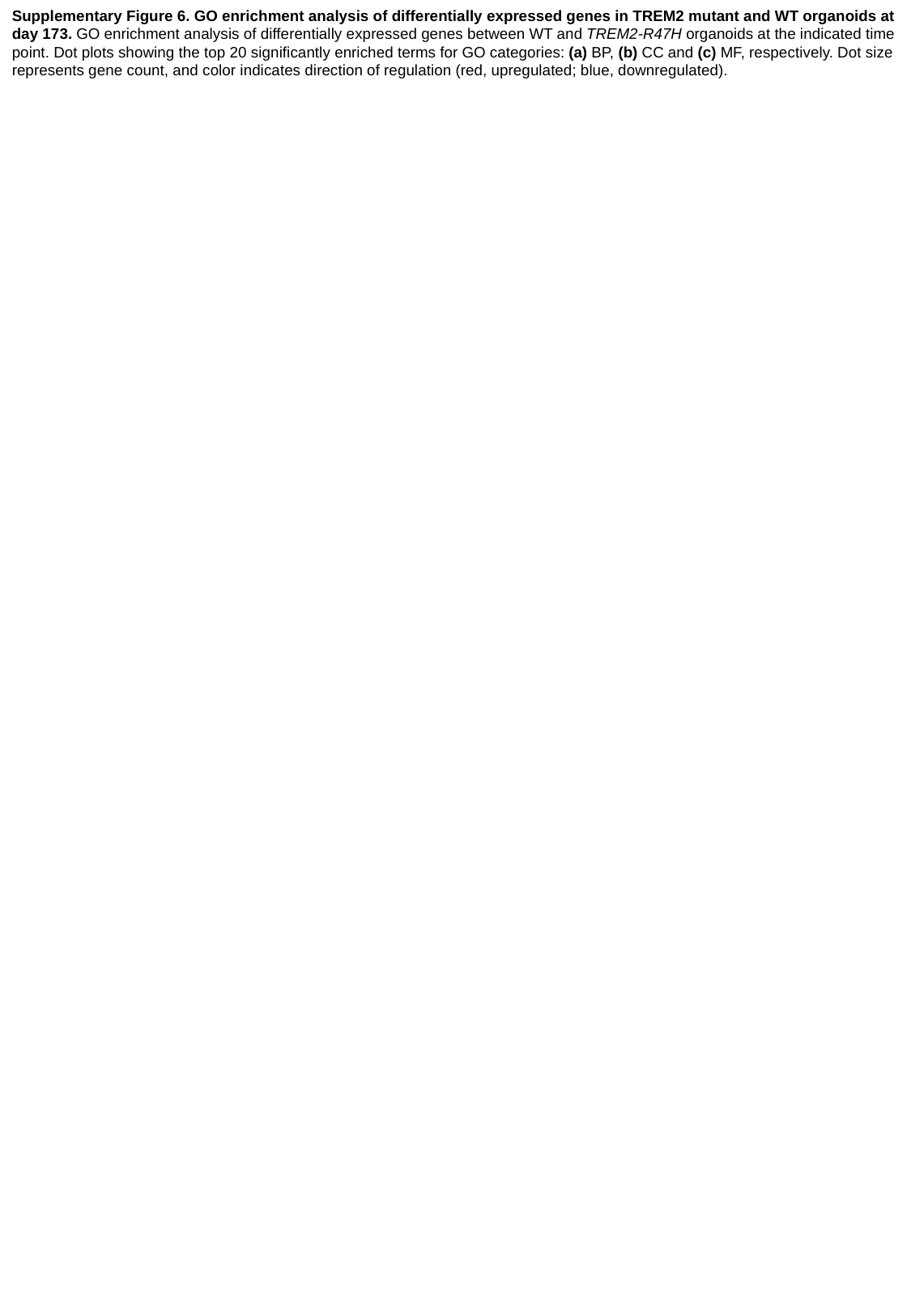

Supplementary Figure 6. GO enrichment analysis of differentially expressed genes in TREM2 mutant and WT organoids at day 173. GO enrichment analysis of differentially expressed genes between WT and TREM2-R47H organoids at the indicated time point. Dot plots showing the top 20 significantly enriched terms for GO categories: (a) BP, (b) CC and (c) MF, respectively. Dot size represents gene count, and color indicates direction of regulation (red, upregulated; blue, downregulated).

### Slide 10
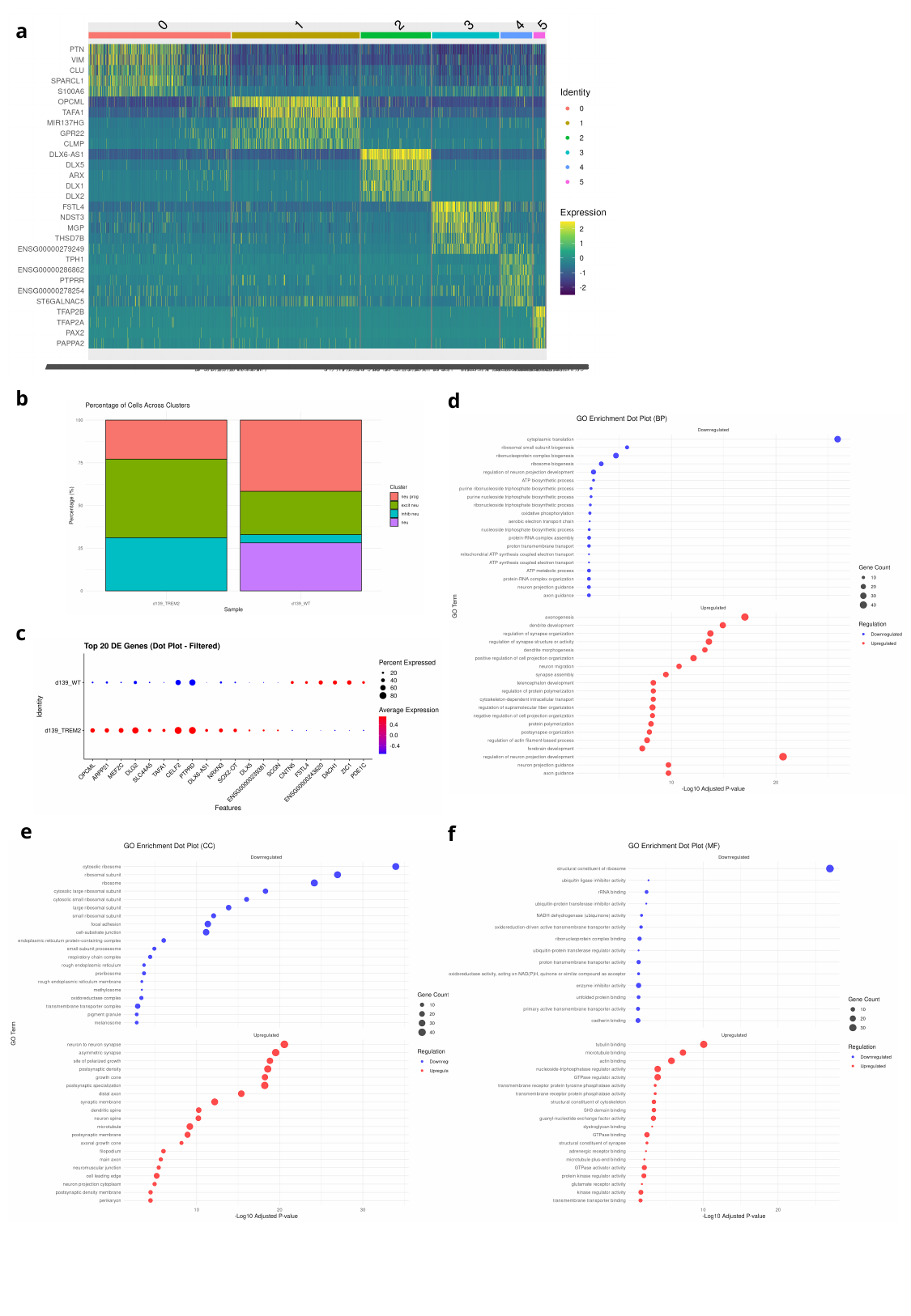

a
b
d
c
e
f

### Slide 11
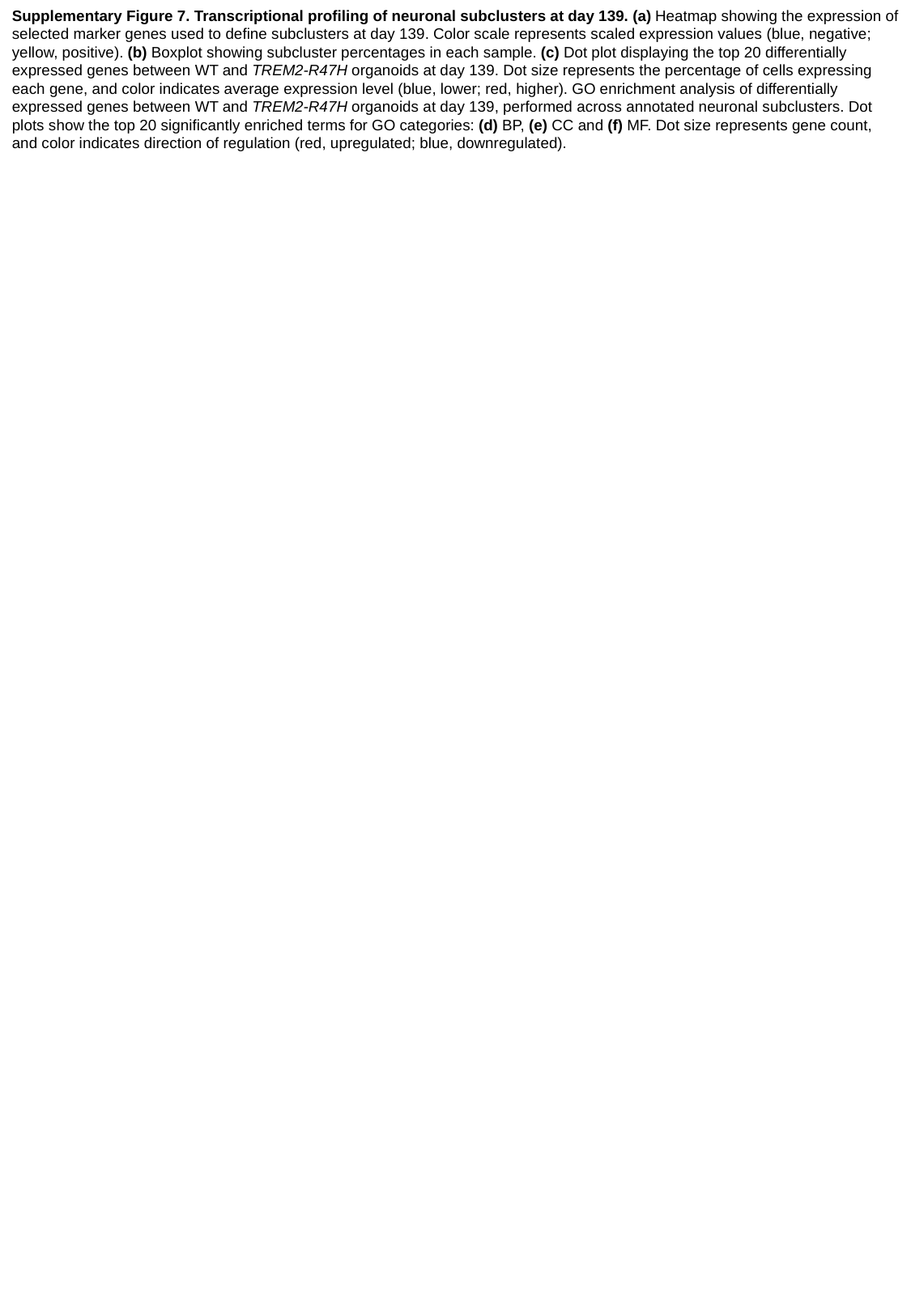

Supplementary Figure 7. Transcriptional profiling of neuronal subclusters at day 139. (a) Heatmap showing the expression of selected marker genes used to define subclusters at day 139. Color scale represents scaled expression values (blue, negative; yellow, positive). (b) Boxplot showing subcluster percentages in each sample. (c) Dot plot displaying the top 20 differentially expressed genes between WT and TREM2-R47H organoids at day 139. Dot size represents the percentage of cells expressing each gene, and color indicates average expression level (blue, lower; red, higher). GO enrichment analysis of differentially expressed genes between WT and TREM2-R47H organoids at day 139, performed across annotated neuronal subclusters. Dot plots show the top 20 significantly enriched terms for GO categories: (d) BP, (e) CC and (f) MF. Dot size represents gene count, and color indicates direction of regulation (red, upregulated; blue, downregulated).

### Slide 12
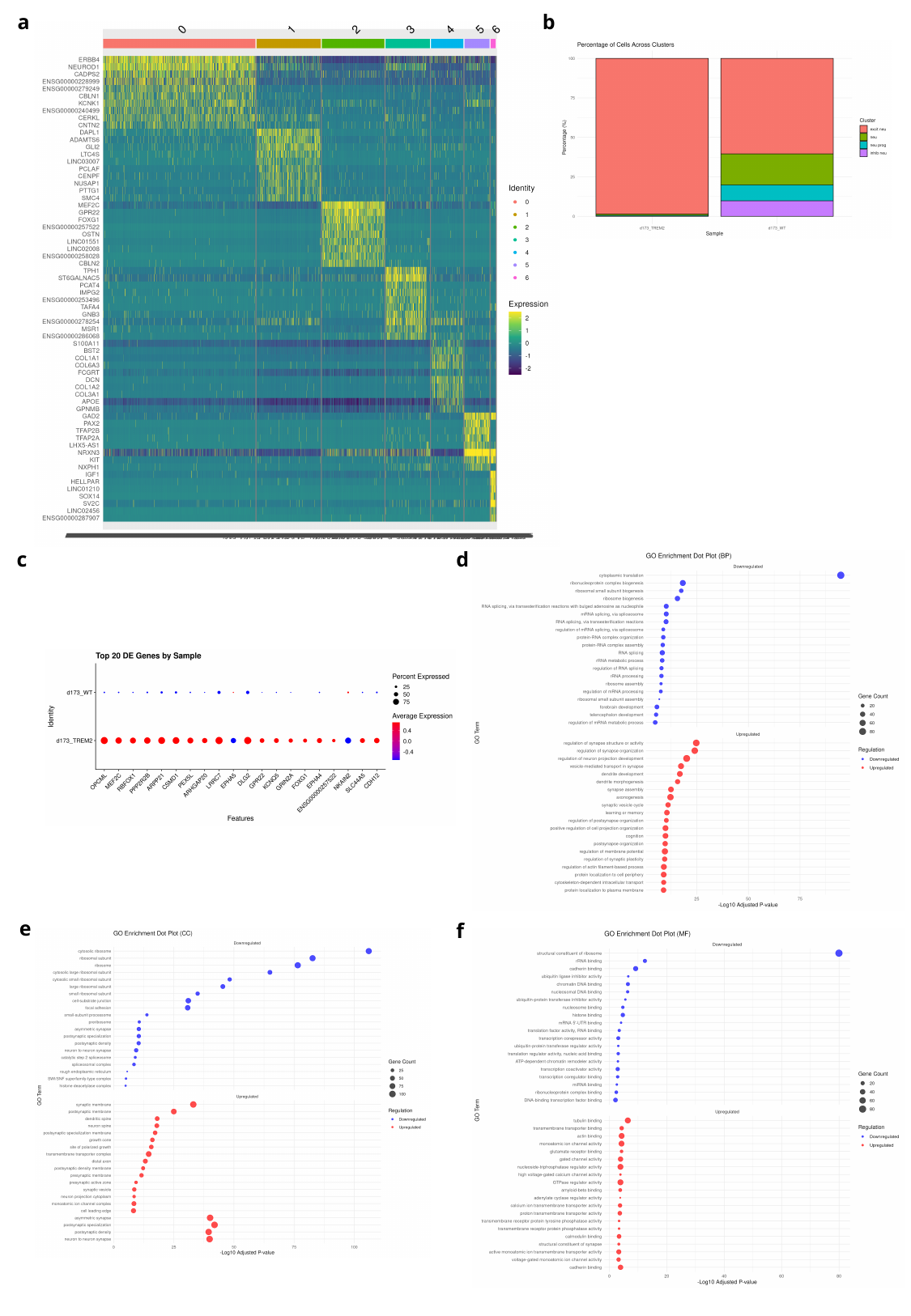

b
a
d
c
e
f

### Slide 13
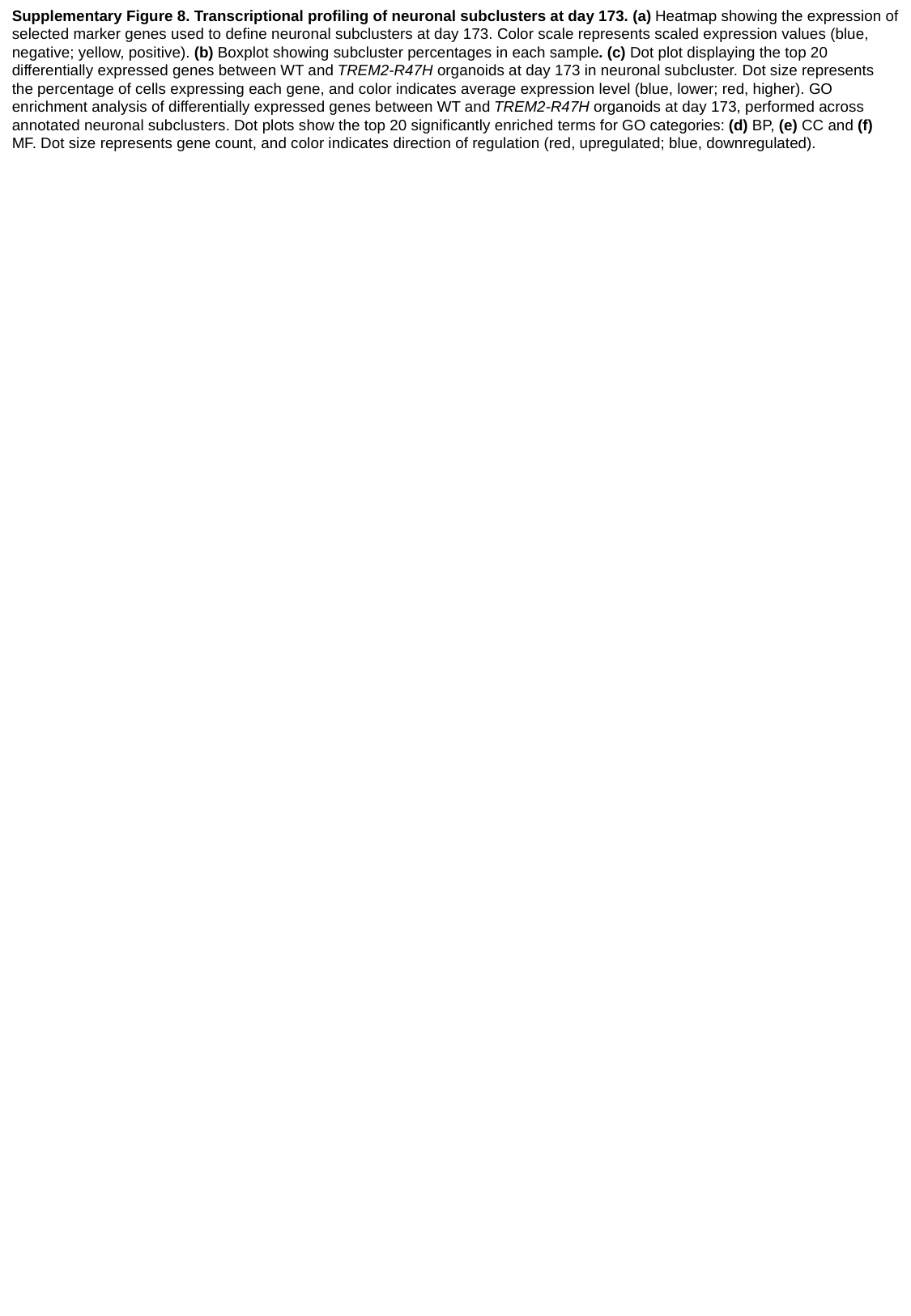

Supplementary Figure 8. Transcriptional profiling of neuronal subclusters at day 173. (a) Heatmap showing the expression of selected marker genes used to define neuronal subclusters at day 173. Color scale represents scaled expression values (blue, negative; yellow, positive). (b) Boxplot showing subcluster percentages in each sample. (c) Dot plot displaying the top 20 differentially expressed genes between WT and TREM2-R47H organoids at day 173 in neuronal subcluster. Dot size represents the percentage of cells expressing each gene, and color indicates average expression level (blue, lower; red, higher). GO enrichment analysis of differentially expressed genes between WT and TREM2-R47H organoids at day 173, performed across annotated neuronal subclusters. Dot plots show the top 20 significantly enriched terms for GO categories: (d) BP, (e) CC and (f) MF. Dot size represents gene count, and color indicates direction of regulation (red, upregulated; blue, downregulated).

### Slide 14
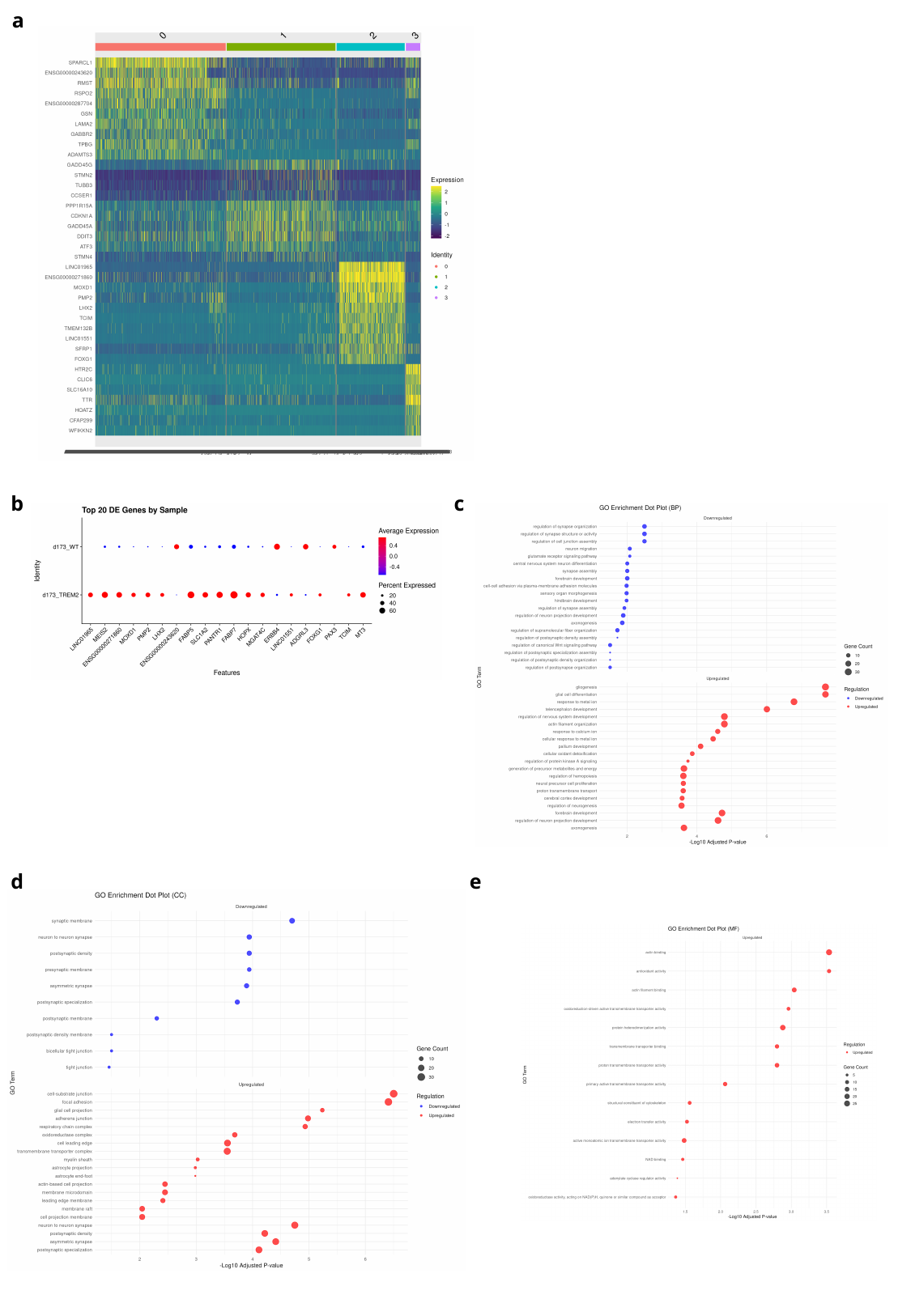

a
c
b
d
e

### Slide 15
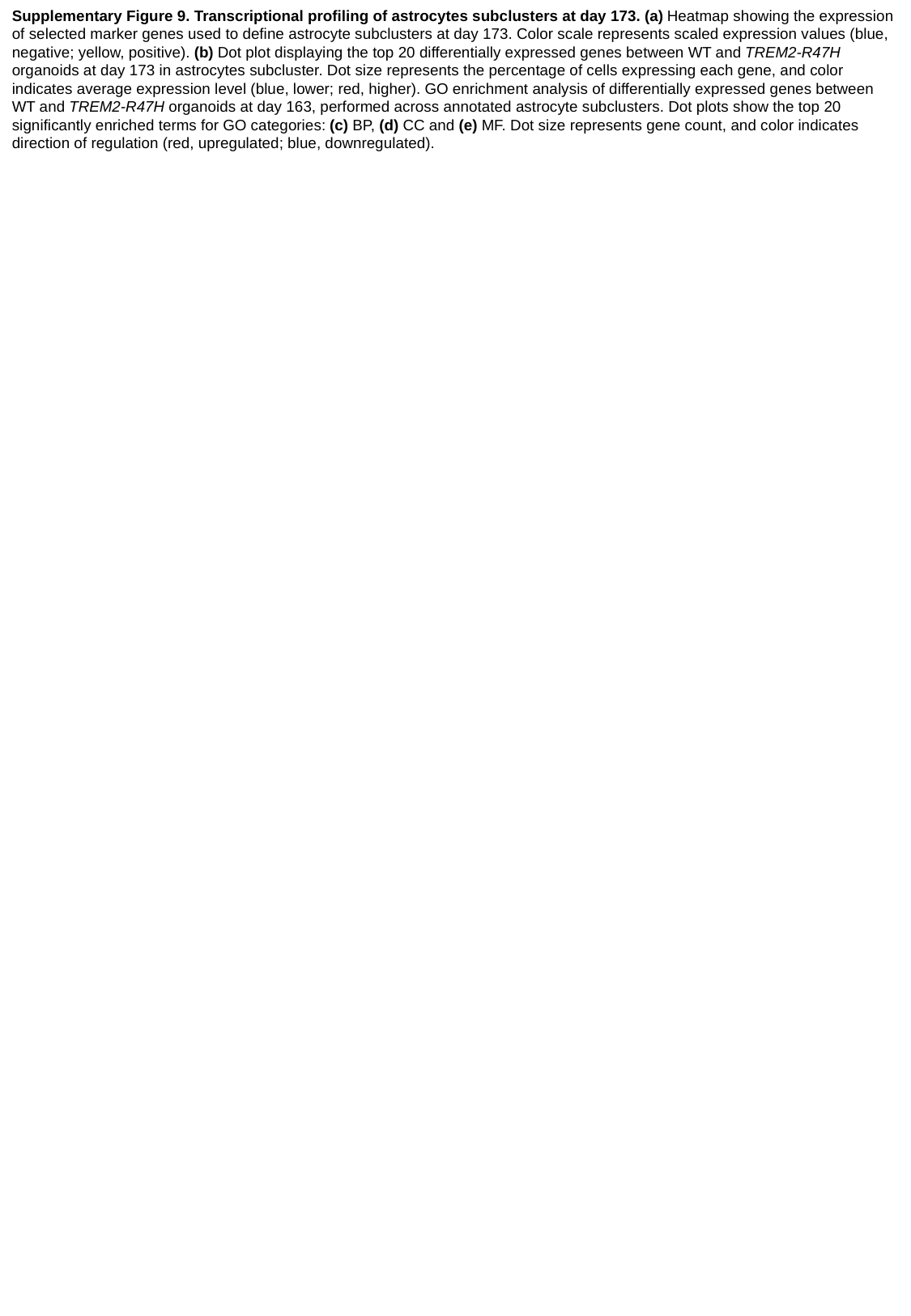

Supplementary Figure 9. Transcriptional profiling of astrocytes subclusters at day 173. (a) Heatmap showing the expression of selected marker genes used to define astrocyte subclusters at day 173. Color scale represents scaled expression values (blue, negative; yellow, positive). (b) Dot plot displaying the top 20 differentially expressed genes between WT and TREM2-R47H organoids at day 173 in astrocytes subcluster. Dot size represents the percentage of cells expressing each gene, and color indicates average expression level (blue, lower; red, higher). GO enrichment analysis of differentially expressed genes between WT and TREM2-R47H organoids at day 163, performed across annotated astrocyte subclusters. Dot plots show the top 20 significantly enriched terms for GO categories: (c) BP, (d) CC and (e) MF. Dot size represents gene count, and color indicates direction of regulation (red, upregulated; blue, downregulated).

### Slide 16
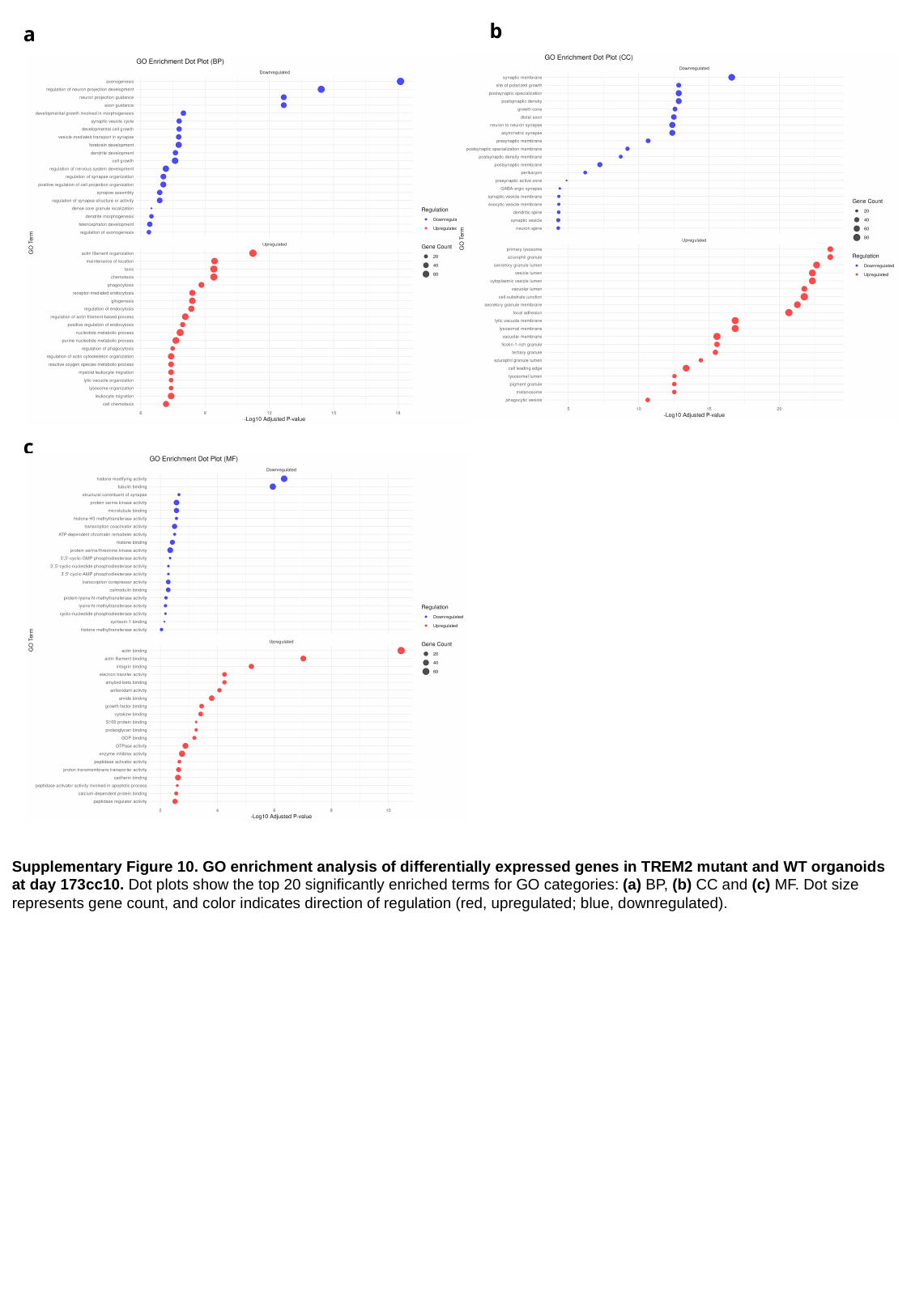

b
a
c
Supplementary Figure 10. GO enrichment analysis of differentially expressed genes in TREM2 mutant and WT organoids at day 173cc10. Dot plots show the top 20 significantly enriched terms for GO categories: (a) BP, (b) CC and (c) MF. Dot size represents gene count, and color indicates direction of regulation (red, upregulated; blue, downregulated).

### Slide 17
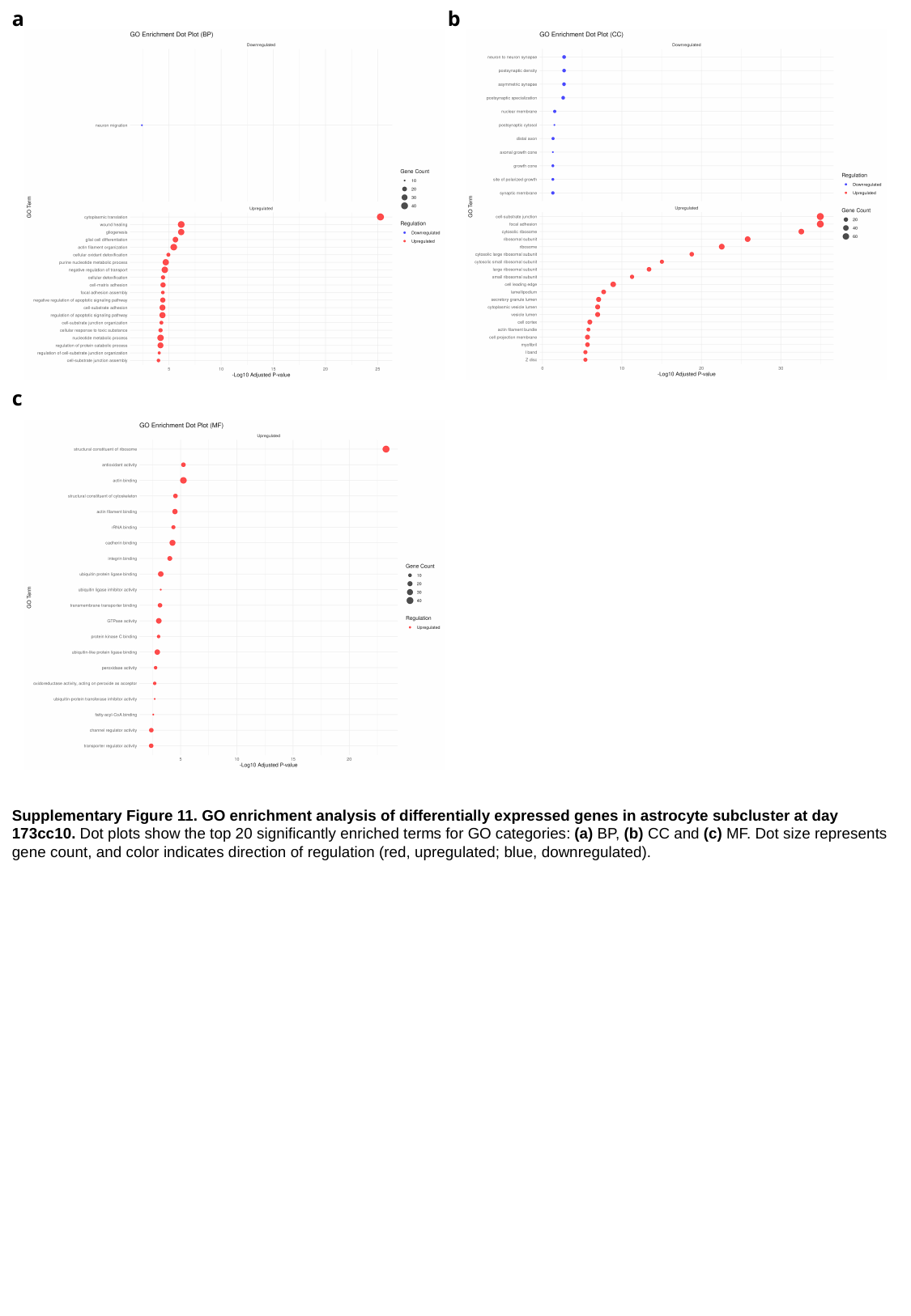

a
b
c
Supplementary Figure 11. GO enrichment analysis of differentially expressed genes in astrocyte subcluster at day 173cc10. Dot plots show the top 20 significantly enriched terms for GO categories: (a) BP, (b) CC and (c) MF. Dot size represents gene count, and color indicates direction of regulation (red, upregulated; blue, downregulated).

### Slide 18
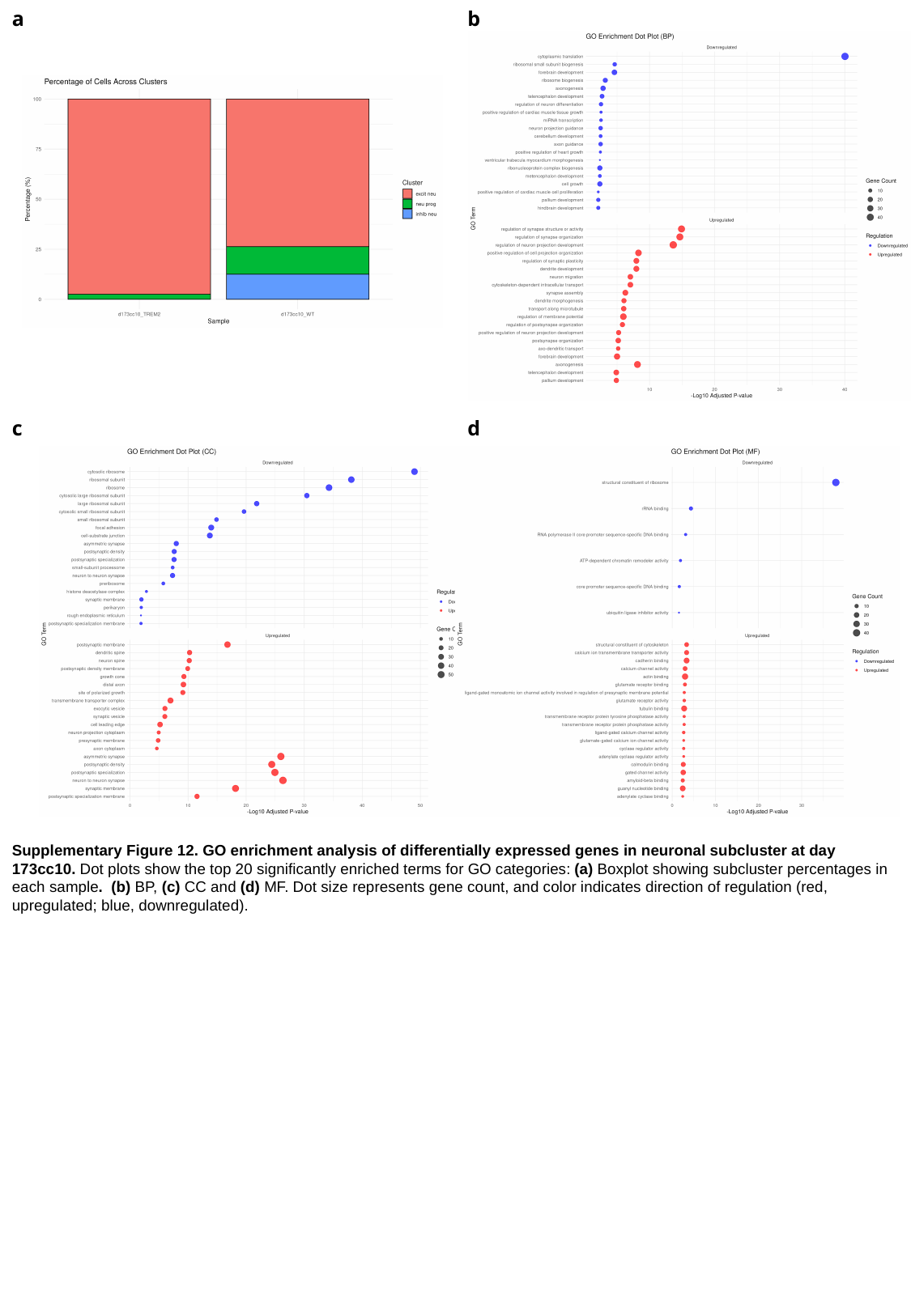

a
b
d
c
Supplementary Figure 12. GO enrichment analysis of differentially expressed genes in neuronal subcluster at day 173cc10. Dot plots show the top 20 significantly enriched terms for GO categories: (a) Boxplot showing subcluster percentages in each sample. (b) BP, (c) CC and (d) MF. Dot size represents gene count, and color indicates direction of regulation (red, upregulated; blue, downregulated).

### Slide 19
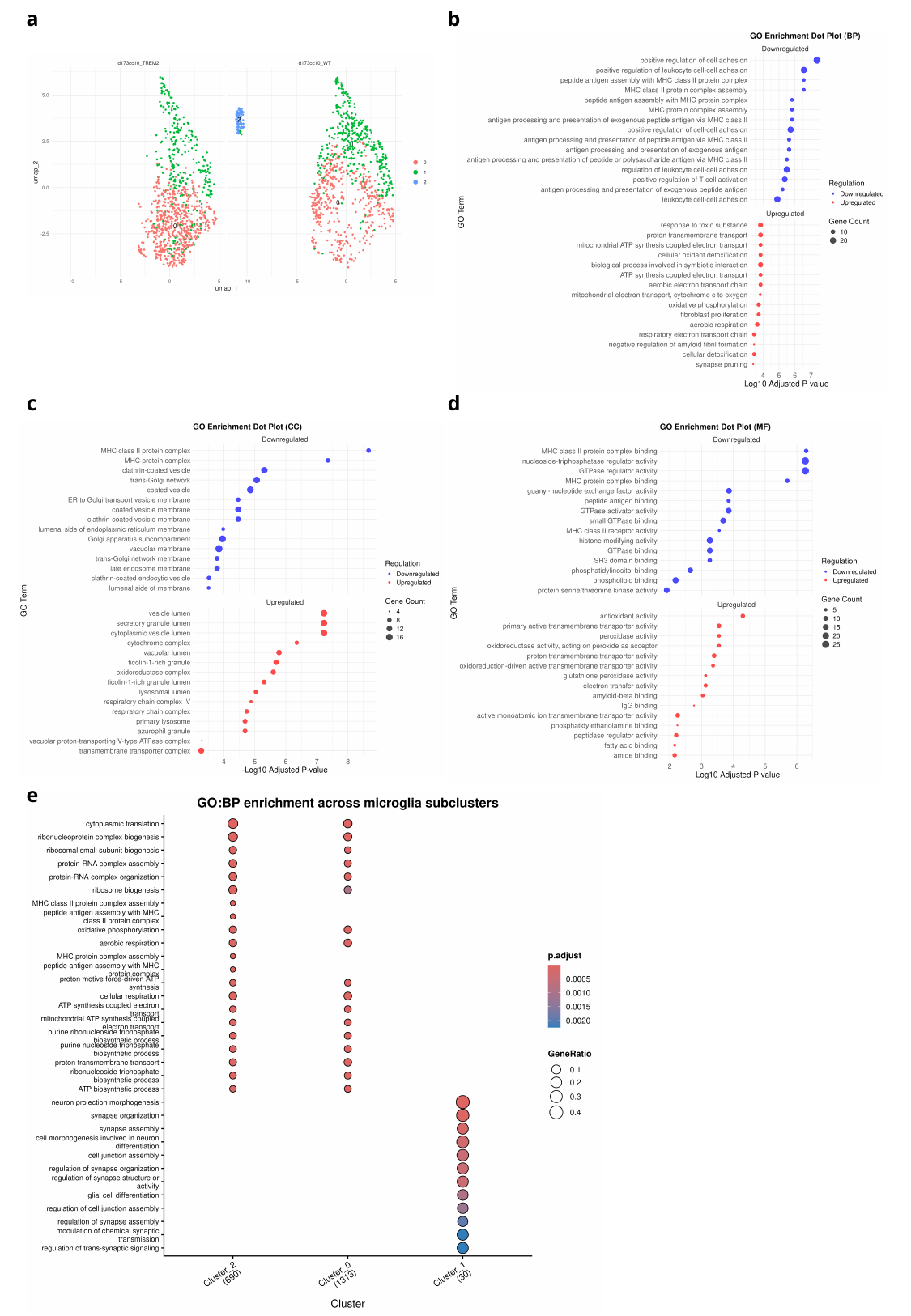

a
b
c
d
e

### Slide 20
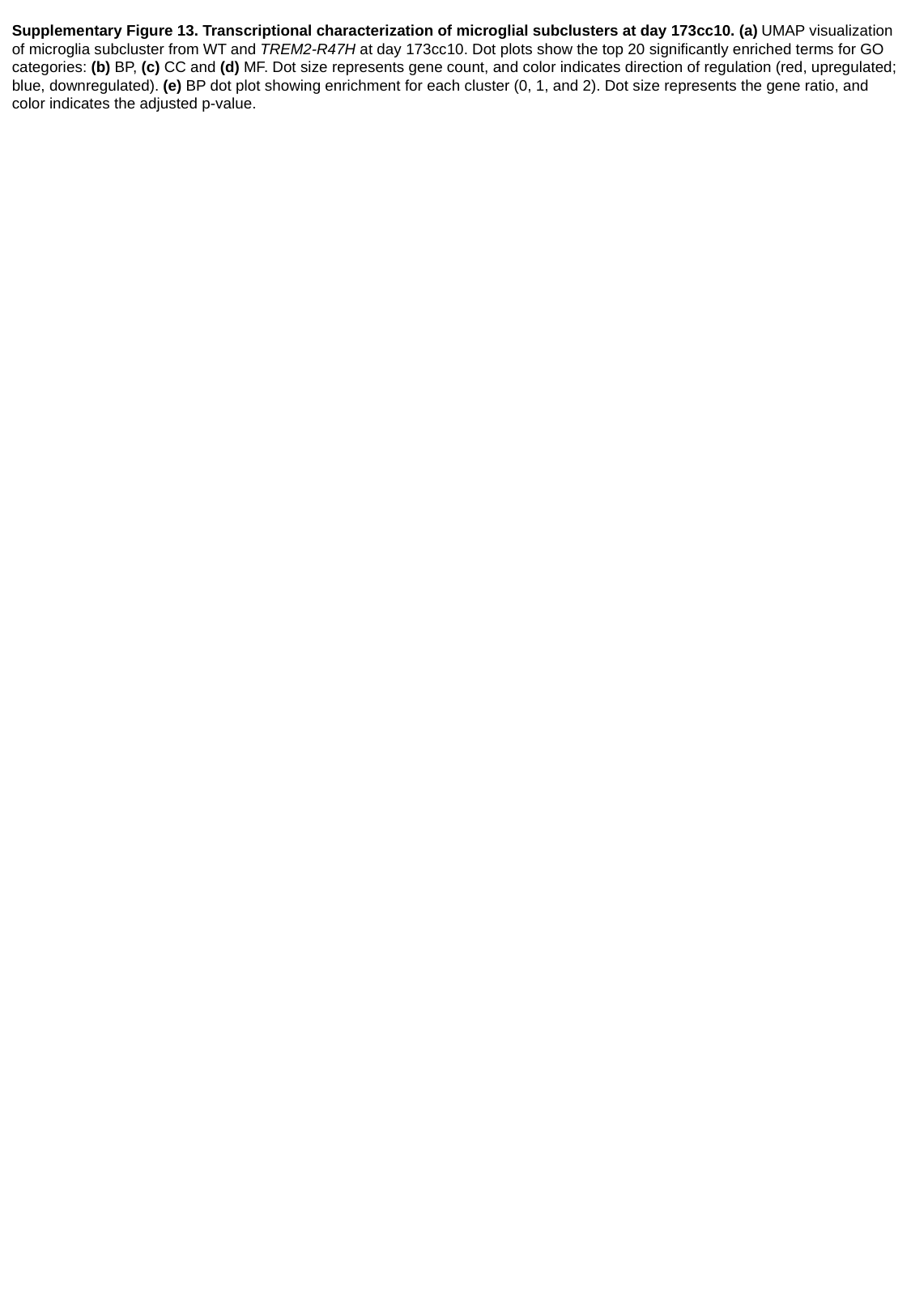

Supplementary Figure 13. Transcriptional characterization of microglial subclusters at day 173cc10. (a) UMAP visualization of microglia subcluster from WT and TREM2-R47H at day 173cc10. Dot plots show the top 20 significantly enriched terms for GO categories: (b) BP, (c) CC and (d) MF. Dot size represents gene count, and color indicates direction of regulation (red, upregulated; blue, downregulated). (e) BP dot plot showing enrichment for each cluster (0, 1, and 2). Dot size represents the gene ratio, and color indicates the adjusted p-value.
